# Functional specialization of human salivary glands and origins of proteins intrinsic to human saliva

**DOI:** 10.1101/2020.02.12.945659

**Authors:** Marie Saitou, Eliza Gaylord, Erica Xu, Alison May, Lubov Neznanova, Sara Nathan, Anissa Grawe, Jolie Chang, William Ryan, Stefan Ruhl, Sarah M. Knox, Omer Gokcumen

**Affiliations:** Department of Biological Sciences, University at Buffalo, The State University of New York; Section of Genetic Medicine, Department of Medicine, University of Chicago; Program in Craniofacial Biology, Department of Cell and Tissue Biology, School of Dentistry, University of California, San Francisco; Currently at Weill-Cornell Medical College, Physiology and Biophysics Department.; Department of Oral Biology, School of Dental Medicine, University at Buffalo, The State University of New York; Department of Otolaryngology, School of Medicine, University of California, San Francisco

## Abstract

Salivary proteins are essential for maintaining health in the oral cavity and proximal digestive tract and serve as a diagnostic window into human disease. However, their precise organ origins remain unclear. Through transcriptomic analysis of major adult and fetal salivary glands, and integration with the saliva proteome and transcriptomes of 28+ organs, we linked human saliva proteins to their source, identified salivary gland-specific genes, and uncovered fetal- and adult-specific gene repertoires. Our results also provide new insights into the degree of gene retention during maturation and suggest that functional diversity between adult gland-types is driven by specific dosage combinations of hundreds of transcriptional regulators rather than a few gland-specific factors. Finally, we demonstrate the hitherto unrecognized heterogeneity of the human acinar cell lineage. Our results pave the way for future investigations into glandular biology and pathology, as well as saliva’s use as a diagnostic fluid.

## INTRODUCTION

The mouth is the main portal of entry to the gastrointestinal tract with saliva serving as the quintessential gatekeeper (Dawes et al., 2015; Ruhl, 2012). Saliva exerts a multitude of important functions in the oral cavity and beyond that depend upon its repertoire of proteins. These functions include the breakdown of dietary starch by the salivary enzyme amylase (Ragunath et al., 2008), the provision of calcium phosphate to maintain mineralization of tooth enamel (Moreno et al., 1979), and host defense against pathogenic microorganisms while maintaining a beneficial commensal microbiome in the mouth (Cross and Ruhl, 2018; Heo et al., 2013). Saliva also possesses physicochemical properties keeping the oral cavity moist and well lubricated, which are equally provided by saliva proteins, especially mucins (Frenkel and Ribbeck, 2015; Tabak et al., 1982). Thus, genetic and dosage variation in the saliva proteome will have important biomedical consequences (Helmerhorst and Oppenheim, 2007; Pajic et al., 2019; Poole et al., 2019; Xu et al., 2017). At the extreme, malfunctioning of the salivary glands due to, for instance, radiation treatment of head and neck cancer, or caused by the relatively common autoimmune disease, Sjögren’s syndrome, can lead to major disruptions in protein homeostasis within saliva causing severe complications in oral health that debilitate patient quality of life (Mavragani and Moutsopoulos, 2019; Vissink et al., 2015). Therefore, understanding how the composition of the saliva proteome is attained and regulated remains an important avenue of inquiry.

A major complication in studying saliva and harnessing its proteome profile for diagnostic applications is the complexity of this oral biofluid due to it being a mixture of components derived from multiple sources. Saliva gets predominantly synthesized and secreted by three major pairs of anatomically and histologically distinct craniofacial secretory organs: the parotid, submandibular, and sublingual salivary glands (**Figure 1A**). Each of these gland types produces a characteristic spectrum of salivary proteins that are thought to be predominantly based on their unique composition of mucous and serous acinar cells. These ’intrinsic’ (*i.e.*, *de-novo* synthesized by the glands) proteins sustain most of the primary functions of the saliva. In addition, saliva also contains ’extrinsic’ proteins that originate in other organs and systems, including the bloodstream and cells lining the oral integuments (Heller et al., 2017; Ruhl, 2012; Yan et al., 2009). Indeed, proteins can diffuse into saliva from other organs and organ systems via crossing capillary barriers, interstitial spaces, and the membranes of acinar and ductal cells. Whole mouth saliva also contains a plethora of proteins derived from the oral microbiome (more than 700 different species of microorganisms cohabit saliva), adding to the functional complexity of human saliva (Lamont et al., 2018; Mark Welch et al., 2016).

**Figure 1.**
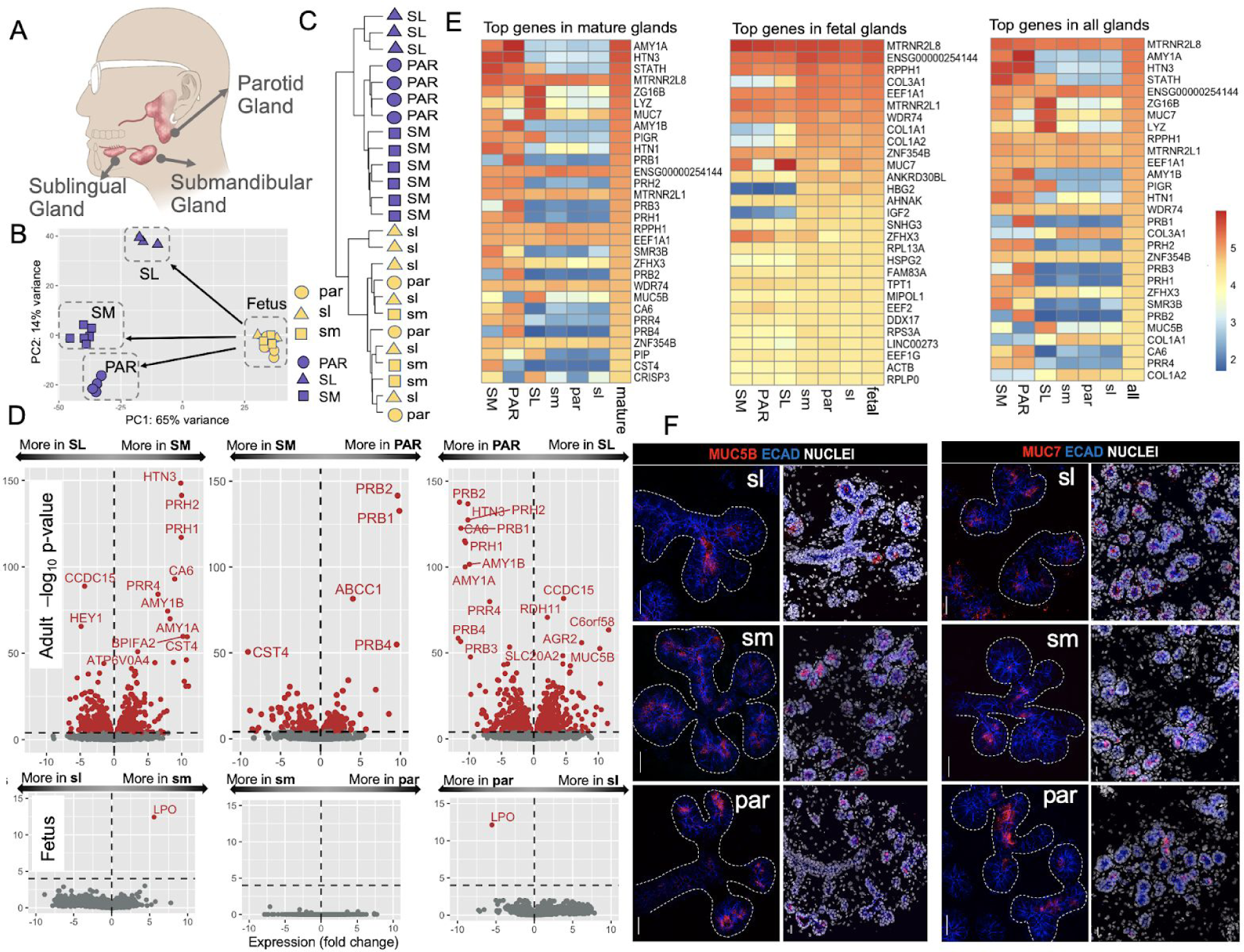
Overview of the transcriptomics analysis. **(A)** Anatomical location of the three major glands in humans. **(B)** Principal component analysis of the gene expression levels in adult and fetal parotid, submandibular, and sublingual glands. Adult gland types show clear separation based on their gene expression without any a priori hypothesis, while fetal glands cannot be separated based on their gene expression alone. Blue symbols represent adult samples and yellow symbols represent fetal samples. Triangle, square, and circle shapes represent parotid, submandibular, and sublingual glands, respectively. **(C)** Hierarchical clustering analysis of adult and fetal parotid, submandibular, and sublingual, glands based on all the transcriptome data without a priori clustering information. **(D)** Transcriptional comparison of adult and fetal major salivary glands. Volcano plots showing the expression differences between glands in a pairwise fashion for adult (top) and fetus (bottom). The x-axis indicates gene expression fold changes. The y-axis indicates -log10 value of adjusted p-value. Genes with significantly different expressions between glands are indicated in red on the plots (adjusted p-value < 0.0001), others were shown in gray. **(E)** Heatmaps of the log10 normalized expression of gland-specific genes. The top 30 expressed genes in the mature, fetal and both types of glands are shown. Note that the top gene, *RN7SL1*, was excluded because of its role as a housekeeping gene. *MUC7* is expressed in both adult and fetal glands and is enriched in adult sublingual glands. *HTN*, *AMY*, and *STATH* are highly expressed in adult parotid and submandibular glands. *CST* genes are expressed in the adult submandibular gland, and PRB genes are expressed in the adult parotid gland, specifically. (**F**) Immunofluorescent localization of *MUC7* and *MUC5B* in fetal glandular tissues shows that both mucins are expressed at the protein level in all fetal glands. Left panel: ECAD = E cadherin; NUC = nuclei. Scale bar = 25 µm.

A multitude of studies has been conducted to catalog salivary proteins and distinguish intrinsic from extrinsic protein components (Grassl et al., 2016), including those that specifically investigated ductal secretions (Denny et al., 2008). These studies include proteomic analyses comparing whole saliva or saliva collected from the ducts of the parotid or submandibular/sublingual (Castagnola et al., 2017; Ogawa et al., 2011) to body fluids such as plasma, urine, cerebrospinal fluid, and amniotic fluid (Loo et al., 2010). However, discrepancies among published data sets, likely due to variation in collection procedures, sample integrity, storage conditions, sample size, and analytical methods (Helmerhorst and Oppenheim, 2007; Robinson et al., 2008; Thomadaki et al., 2011), have impaired the establishment of a robust catalog of proteins – an outcome that has so far severely hampered the use of saliva as a physiological and pathophysiological research tool and as a reliable fluid for disease diagnosis.

To address this gap in knowledge, a major effort has been undertaken in compiling a systematic knowledge database of proteins in human saliva, the recently established Human Salivary Proteome Wiki (https://salivaryproteome.nidcr.nih.gov/), a community-based, collaborative web portal that harbors information on salivary proteins identified by high-throughput proteomic technologies including mass spectrometry. The accumulating data on the protein content of saliva led to several intriguing questions in regard to human saliva: *What are the relative contributions of different salivary glands to the composition of the salivary proteome? How different are the secretomes of the three major adult human salivary gland types and what genes drive these differences? What secretory and non-secretory proteins are uniquely produced by the salivary glands? And finally, what can we learn by studying fetal salivary glands to define the adult secretory function?* To help answer these questions, we have constructed the first comprehensive dataset of the transcriptomes of the major human salivary glands types and integrated these data with the mass spectrometric proteomic analyses from the Human Salivary Proteome Wiki and Human Protein Atlas (Uhlen et al., 2010). We sequenced total RNA of 25 salivary gland samples collected from adult and fetal human submandibular, sublingual, and parotid glands and analyzed transcriptomes across gland type and developmental stage, and further compared these to the transcriptomes of 52 organ systems (GTEx database).

## RESULTS AND DISCUSSION

### Distinct transcript profiles between adult but not fetal salivary gland types indicate that functional specialization occurs during late-stage development

To comprehensively identify gene expression differences between the three major salivary gland types, we conducted a transcriptome analysis of multiple healthy male and female adult and fetal sublingual (SL, sl), submandibular (SM, sm), and parotid (PAR, par) gland tissues (Table S1, **Figure 1A**). Human salivary gland development begins at 6-8 weeks, with the formation of a branched structure with clearly defined end buds (pre-acini) by 16 weeks and lumenized acini by 20 weeks (Kumagai and Sato, 2003). The period after 20 weeks is associated with cytodifferentiation and the presence of intercalated/striated ducts and is characterized as the last stage of salivary gland development (Ianez et al., 2010). Salivary glands are considered fully differentiated by 28 weeks, as noted by the presence of secretory vesicles and the expression of secretory protein BPIFA1/SPLUNC1(Zhou et al., 2006). Here we utilized glandular tissues taken at 22-23 weeks of age and, based on these previous studies, define this age group as late-stage development.

We comparatively analyzed the expression levels of 167,278 transcripts consolidated into 40,882 coding and noncoding genes using a DeSeq2 pipeline (Love et al., 2014) and gene-transcript information on Ensembl (http://asia.ensembl.org/index.html) (**Table S2, Supplementary Methods**). We were able to clearly differentiate mature glands from fetal glands without any *a priori* hypothesis based only on the first principal component of transcriptome data (**Figure 1B**). The second principal component of the transcriptome data evidently separated the mature gland types. However, the same analysis could not differentiate among fetal gland types. We verified these results using a hierarchical clustering analysis (**Figure 1C**), where the transcripts of the mature glands significantly clustered into a major branch distinct from that of the fetal glands. Moreover, we found that the transcripts of the mature glands further branched up according to their glandular origin, whereas those of the fetal glands did not. We were able to quantify these observations using a Pearson correlation analysis (**Figure S1**). As expected for tissues composed of similar cell types, the adult PAR gland exhibited greater similarity in global gene expression to the SM than the SL gland.

To determine which transcripts account for the differences between the gland types, we conducted a comparative analysis between glandular transcripts (**Table S2**, **Figure 1D**). As expected from the global analysis described above, we found hundreds of transcripts that are differentially expressed among mature glands. Not surprisingly, gene ontology (GO)analysis showed genes found predominantly expressed in the fetal tissues to be significantly enriched in categories linked to growth and development, including the cell cycle, cell division, and other fundamental cellular processes (**Table S3)**. Consistent with the functional maturation of the tissues, we found that differences among mature gland types were mainly due to genes that code for secreted proteins, as defined by the Human Protein Atlas (https://www.proteinatlas.org/)).

Indeed, we confirmed the gland-type specific and abundant expression of a number of genes coding for secreted proteins routinely found in saliva (Ruhl, 2012) (**Figure 1E**) (Ruhl, 2012). These included genes that were shared by two gland types but not by the third or that were expressed exclusively by one gland type. For example, the genes for *AMY1*, salivary histatins (*HTN1, HTN3*), and salivary acidic proline-rich proteins (*e.g.*, *PRH1, PRH2*) were expressed at highest levels in both PAR and SM gland but much less or not at all in SL tissue. Transcripts for *MUC7* were virtually absent in PAR tissue but were expressed at highest levels in the SL and to a lesser extent in the SM (Nielsen et al., 1996) whereas transcript levels for salivary cystatins (*e.g.*, *CST1*, *CST2 (Dickinson et al., 2002)*) were found at highest levels in SM, but not in SL tissue. In addition to these, we found that genes previously believed to be expressed by both SM and PAR glands were exclusively expressed by one but not the other. These included mRNA encoding members of the basic proline-rich protein BstNI subfamily (*PRB1*, *2*, and *4*) and cystatin family (*CST4*), the expression of which was found restricted to the PAR and SM, respectively.

We also identified a substantial number of genes coding for secretory proteins that have not been previously described to differ between gland types (**Table S2**). For example, we found kallikrein 1 (*KLK1*), low-density lipoprotein receptor-related protein 1B (*LRP1B)*, mucin-like 1 (*MUCL1/SBEM (Miksicek et al., 2002)*), carbonic anhydrase (*CA6 (Parkkila et al., 1990)*) and *C6orf46/SSSP1* (skin and saliva secreted protein 1; cell origin of protein is unknown (Gerber et al., 2013)) were expressed by the PAR and SM and absent from the SL while contactin 5 (*CNTN5*) and secreted phosphoprotein 1/osteopontin (*SPP1*) were restricted to the PAR and SL glands, respectively. Similarly, *CRISP3,* a gene identified as a novel early response gene that may participate in the pathophysiology of the autoimmune lesions of Sjogren’s disease (Tapinos et al., 2002) and has previously been shown to be expressed by the acinar cells of human labial glands (Laine et al., 2007), was highly expressed by the SL and to a lesser extent by the SM, but absent from the PAR. Moreover, based on previous proteomics work reported in Human Salivary Proteome Wiki, these proteins are found in the whole saliva and, thus, they are exciting targets for future research into salivary function, diagnostic and health.

A small fraction of secreted genes were also found to be exclusively expressed by only one adult gland type. For example, transcripts for low-density lipoprotein receptor-related protein 2 (*LRP2/*megalin), a multiligand uptake receptor that is involved in protein reabsorption (Christensen and Birn, 2002), were only found in PAR tissue; endothelin 3 (*EDN3 (Gurusankar et al., 2015)*) was highly enriched in the SM; and the mucous components *FCGBP* (Pelaseyed et al., 2014), *AGR2* (Park et al., 2009), and trefoil factor 1 (*TFF1* (Chaiyarit et al., 2012)) along with a gene of unknown function enriched in mucous tissues, *C6orf58/LEG1 (Pelaseyed et al., 2014)*, were restricted to the SL. Some of the proteins that these genes encode are found in saliva (e.g., C6orf58/LEG1 (Ramachandran et al., 2006)) while others have a negligible presence (e.g., LRP2/megalin), although whether this deficiency is due to protein degradation or if they are simply not secreted into the ductal lumens is yet to be determined.

Adult gland types also differed significantly in the expression of sets of immune-related secretory genes. For instance, through GO analysis (**Table S3**) we found distinct complement cascades and immunoglobulin production pathways (*e.g.*, IGHV1-58 and C6) that were shared by the PAR and SM but were different from those shared by the SM and SL. Indeed, the SM and SL showed substantially greater levels of transcripts for genes that are primarily responsible for acquired and innate immune functions in the whole saliva, e.g., secretory immunoglobulin (S-IgA), IgG. Also, other proteins involved in host defense in a broader sense (Fábián et al., 2012) including lysozyme, BPI, BPI-like and PLUNC proteins, cystatins, mucins, peroxidases, statherin, and others. This observation raises the possibility that SM and SL may provide a background state of innate immunity in the oral cavity in the absence of food or pathogenic stimuli. Although it is not known whether these differences in immune response gene expression between glands result in tissue-specific outcomes in humans, glandular inflammatory conditions (sialadenitis) show a predilection for certain gland types. For example, Heerfordt syndrome causes parotitis (Takahashi and Horie, 2002), whereas chronic sclerosing sialadenitis predominantly affects the SM glands (Gupta et al., 2015).

Not surprisingly, the proportion of total gene transcripts that encode secreted proteins was significantly higher in mature glands than in their fetal counterparts (p < 0.05, Mann-Whitney test, **Figure S2**). However, a number of secretory genes present in saliva were also expressed at significant levels in fetal glands, albeit lower than found in adult glands (**Figure 1E and 1F**). These included *HTN1*, *MUC7, MUC5B, CRISP3, ZG16B,* and *LPO.* Although genes such as *LPO* and *CRISP3* showed specific enrichment of transcripts in gland types that matched the adult tissues, i.e., *LPO* was enriched in fetal and adult parotid and submandibular and *CRISP3* in fetal and adult submandibular and sublingual glands; surprisingly, many of these genes did not match the tissue-specific expression patterns of the adult organs. For example, transcripts for *MUC7* and *MUC5B*, which are expressed exclusively by the adult submandibular and sublingual glands, were expressed by all fetal gland types including the par (**Figure 1E)**. Such an outcome hints at the possibility of as of yet unknown functions of these genes during fetal development that, in the case of *MUC7* and *MUC5B*, are not required at adult stages.

In addition to the secretory genes, we identified sets of non-secretory genes that also display gland-specific patterns of expression. For example, the adult SL was enriched in transcripts for retinol dehydrogenase 11 (*RDH11)* compared to the PAR and SM glands while transcripts encoding enzymes such as the dopamine degrading monoamine oxidase B (*MAOB*), transcription factors *FEZF2* (a regulator of cell differentiation (Takaba et al., 2015; Zhang et al., 2014)) and *LIM1XB* (LIM homeobox transcription factor 1 beta), co-transporter SLC5A5 and growth factor or steroid receptors such as DNER (Delta and Notch-like epidermal growth factor-related receptor) and progesterone receptor (*PGR*) were almost exclusively expressed by the SM and PAR glands. The multi-drug resistance gene *ABCC1* was highly enriched in the PAR, *GALNT13*, an initiator of O-linked glycosylation of mucins was enriched in the SM and the transcription factor *NKX2-3,* which is required for murine sublingual gland development (Biben et al., 2004), was almost exclusive to the SL. Notably, we found the SM gland to possess few genes showing exclusive expression, consistent with the cell types of this tissue shared with the other glands.

### There is extensive retention of gene transcripts from fetal to adult stages, with the sublingual gland exhibiting the greatest retention of fetal genes

Differences in gene expression between human adult and fetal epithelial organs is poorly understood, with little to no understanding of the genes that are retained or depleted during human salivary gland development or if specific adult gland types show preferential enrichment for fetal genes. To gain insight, we analyzed our RNAseq data sets for genes that were retained or depleted during maturation of the salivary gland types. Overall, we found 8,017 genes were globally expressed at similar levels at both fetal and adult stages (**Figure 2A**), including MTRNR2L8 and WDR74 (also see **Figure 1E**). These genes are enriched for functions related to organ development and adult homeostasis and physiology (**Table S3**). Surprisingly, we also identified factors that had been found to be reduced in expression during murine salivary gland formation, and are known to promote salivary gland development in mice, *e.g.*, fibroblast growth factors 1, 7 and 10 (Mattingly et al., 2015), to be highly retained during maturation of human salivary glands, suggesting species variation in gene retention.

**Figure 2.**
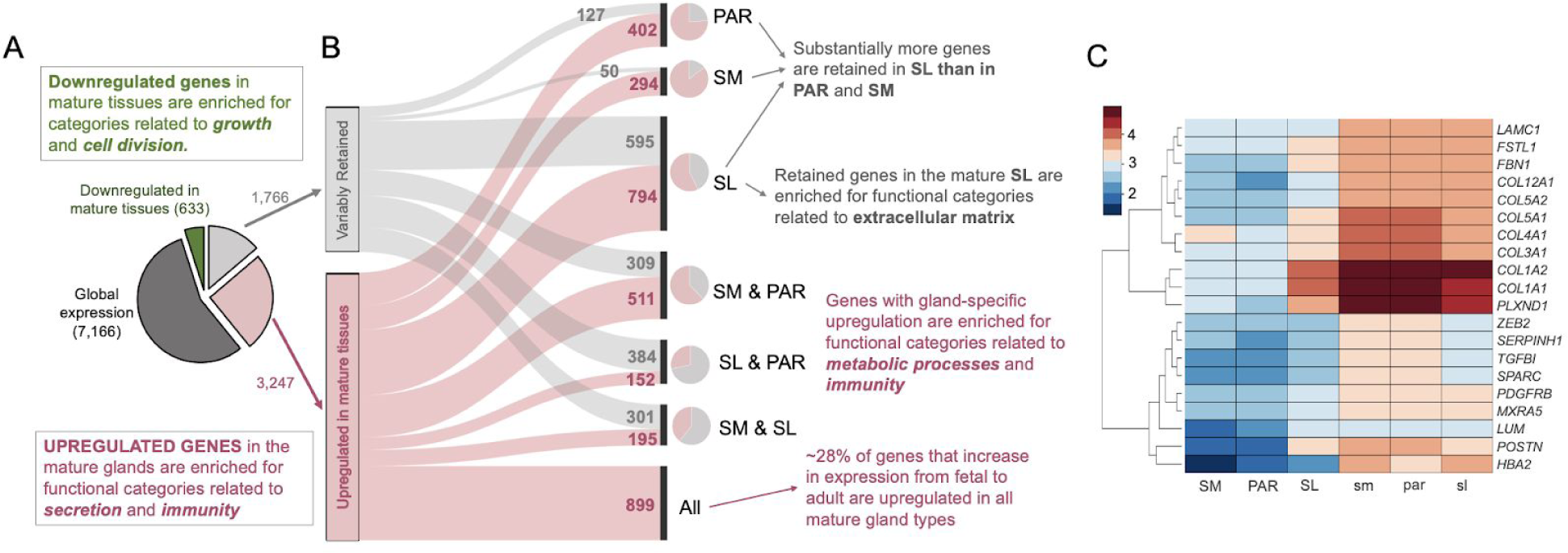
Categorization of genes based on their expression trends in fetal and mature salivary glands. **(A)** The pie-chart on the left indicates the proportion and numbers of genes that we found to be globally expressed i.e., expressed >100 normalized gene count in DESeq2 (Love et al., 2014) and show no significant differences between fetal and mature gland tissues p>0.001 (dark gray) and downregulated in all mature glands as compared to their fetal counterparts (p<0.001, green). The light gray and rose-colored segments indicate genes with expression retained and upregulated in mature salivary glands as compared to their fetal counterparts, respectively. Note that virtually no genes show differential expression between fetal glands with only a few exceptions (see Figure 1D). **(B)** In the middle section, we show a parallel set graph to summarize the breakdown of genes that show variable gene expression in adult glands. This panel indicates how the differential transcriptome repertoires of mature glands are a product of gland-specific retention and upregulation of gene expression. To highlight the different proportions of retained and upregulated genes, we provided pie-charts at the right side of the parallel set graph, where the proportion of retained and upregulated genes for each mature gland is indicated. (**C)** Heatmap showing log_10_ normalized expression of top genes (i.e., with the highest expression in fetal glands, >1000 TPM), the expression of which is retained in only one mature gland from its fetal counterpart but reduced significantly in the other two adult glands. Note that all of these genes are retained in the SL.

Despite extensive similarities in gene retention between adult glands, however, the SL stands out as it retains an additional 1,225 genes from fetal to adult stages (**Figure 2B**), while these genes have reduced expression in the SM and PAR. The genes retained in the SL with the highest expression level are related to extracellular matrix formation and function (**Table S3**, **Figure 2C**). Indeed, some genes including those coding for collagen 1 and 3 isoforms (e.g., *COL3A1*), as well as *SPARC*/osteonectin were retained at fetal-like transcript levels (ten- to hundred-fold higher than in SM and PAR). These results clearly suggest that the SL retains a more fetal-like extracellular matrix that may guide stem cell-mediated repair as has been suggested in other organ systems (Niklason, 2018). For example, periostin (*POSTN*) which is highly retained in the SL has been implicated in stem cell regulation in multiple tissues including bone (Niklason, 2018), heart (Hudson and Porrello, 2017), pancreas (Hausmann et al., 2016), and tendon (Noack et al., 2014).

We also identified a number of highly abundant genes not related to the extracellular matrix that was retained in the adult SL compared to the adult PAR and SM. These included the extracellular glycoprotein Fst-SPARC family member follistatin-like 1 (*FSTL1*), which is an essential regulator of tracheal formation and lung epithelial cell maturation (Geng et al., 2011), and the receptor for semaphorin class 3 ligands plexin D1 (*PLXND1*), which has multiple roles during development e.g., synaptogenesis, heart formation, and vasculogenesis, and is heavily associated with Moebius Syndrome, a developmental neurological disorder that is characterized by paralysis of the facial nerves and variable other congenital anomalies (Tomas-Roca et al., 2015). In regard to Moebius Syndrome, patients show salivary gland dysfunction (Martins Mussi et al., 2016), although whether the tissues themselves are impacted at the morphological levels is unknown.

Intriguingly, despite the extensive changes in cell state that occur during maturation, with the tissue moving from a non-secretory to a secretory organ, very few fetal-expressed genes showed a global loss in gene expression upon transitioning to the adult stage (i.e., absent from all 3 adult glands). Those included genes that are involved in fetal blood e.g., hemoglobin gamma A (*HBGA*), embryonic development e.g., insulin-like growth factor 2 (*IGF2*), and cell proliferation e.g., topoisomerase (DNA) II alpha (TOP2A) as well as a number of transcription factors known to regulate developmental processes in other organ systems e.g., *SOX11* (Huang et al., 2016). *HBGA* is a fetal globin gene, and its low transcript level in adult glands (∼20 transcripts in adults compared to ∼2500 in fetal tissue) ensures the rigor of our study. Thus, our data strongly suggest that fetal genes are highly retained during development, likely having developmental and homeostatic roles. The functional relevance of these retained fetal genes in adult tissue remains yet to be determined.

### The diverse transcription factor (TF) repertoire of mature salivary glands may shape hotspots of hundreds of genes with salivary gland-specific expression across the genome

Given the differences in gene expression between gland types and stages of development, we next tested the hypothesis that TFs function as fundamental players in shaping the overall transcriptome variation among the glandular tissues and, as such, will display specific inter-glandular expression patterns. To address this, we investigated the expression patterns of hundreds of TFs, documenting specific repertoires for each type of salivary gland at fetal and adult stages. We focused on TFs that were either (i) highly abundant in each gland type at each developmental stage, (ii) had salivary gland specific expression, or (iii) had been previously implicated in salivary gland development, disease or cancer (**Figure 3A, Table S4**). More than 60% of known TFs (1025 out of 1648) were expressed (>100 normalized gene count from DESeq) by at least one of the salivary glands during development and maturation, with 661 (∼64%) being expressed in each of the fetal and mature glands suggestive of conserved function during maturation and homeostasis.

**Figure 3:**
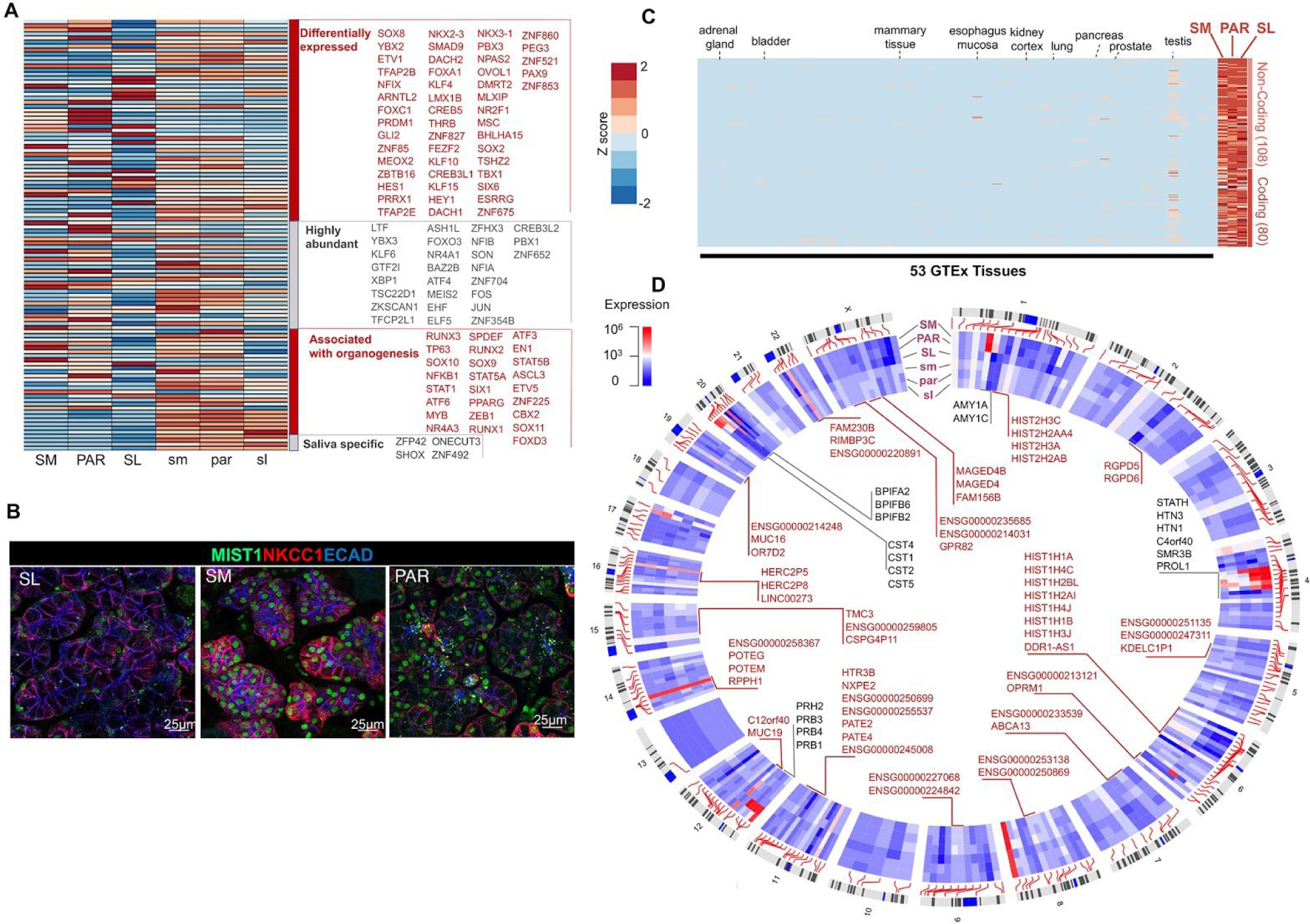
The diverse transcription factor repertoire of mature salivary glands may shape hotspots of salivary gland-specific expression across the genome. A. Heatmap of transcription factor gene (as listed in the TF2DNA database (Pujato et al., 2014)) expression levels across fetal and mature salivary gland tissues. Four categories of transcription factors are shown in the heatmap: Transcription factors that are (i) <u>differentially expressed</u> (P<0.0001) among mature glands, (ii) <u>abundant</u> (>2000, normalized gene count in DESeq) in adult or fetal glands, (iii) previously been <u>associated with organogenesis</u>, and (iv) salivary-gland-specific (that is, >100 normalized gene count from DESeq in the salivary glands but show negligible expression in all 53 GTEx tissue). The colors in the heatmap indicate a positive or negative deviation from the mean for each row. Note that LTF, a known secretory protein in saliva, is listed here due to one of its isoforms, delta lactoferrin, displaying transcription factor activity (He and Furmanski, 1995; Mariller et al., 2007). B. Immunofluorescent analysis of transcription factor BHLHA15/MIST1 in adult glandular tissues. The SM and PAR cells are highly enriched for MIST1 as compared to the SL. ECAD = E cadherin; NKCC1/SLC12A2 = Na-K-Cl cotransporter 1. Scale bar = 25 μm. C. Heatmap of expression levels of genes that have an observable expression in mature salivary glands (>100) but have a negligible expression in other tissues and organs (sum of expression in all 53 GTEx tissues <10 TPM). The specific tissues GTEx database used in this analysis are listed in Table S5. Some relevant epithelial/secretory tissues/organs important for comparison to salivary glands are indicated on top of the heatmap. Deviation from the mean expression for each column is shown as a z-score with a similar scale used in panel A. D. Circos plot showing the locations of genes with salivary gland-specific expression. These genes have a considerable expression in the salivary glands (>100) but show negligible expression in all 53 GTeX tissues. The clusters of genes within 1 Mb of each other are highlighted, and genes that have not previously been reported within the context of salivary glands are indicated in red.

Our analysis identified a host of TFs previously shown through genetic deletion studies in mice to be essential regulators of salivary gland development to be also universally expressed in the developing human glands. These include regulators of acinar cell development (e.g., *SOX2, 9* and *10* (Athwal et al., 2019; Chatzeli et al., 2017; Emmerson et al., 2017)), targets of FGF10 signaling (e.g., *ETV5*), regulators of duct formation (*TFCP2L1* (Yamaguchi et al., 2006) and *YAP1* (Szymaniak et al., 2017)) and of basal stem cells (*TP63* (Song et al., 2018)), and a TF recently discovered to promote salivary organoid initiation from mouse embryonic stem cells (*FOXC1 (Tanaka et al., 2018)*). Notably, a group of TFs including *FOXD3, CBX2,* and *SOX11* were found to be exclusive to the fetal tissue, indicative of developmental rather than homeostatic or physiological roles. This particular group is of high interest to those studying organ bioengineering, wound repair and cancer, as multiple markers present in fetal tissue re-emerge in a variety of cancers (e.g., SOX11 (Yang et al., 2019) and are required for the de-novo generation and regeneration of tissues (Miao et al., 2019; Sock et al., 2004), but their identity remains unclear due to the absence of information on the fetal organs.

Despite the extensive retention of TF expression between fetal and adult stages, TF transcript levels do vary among gland types and between fetal and mature stages (**Figure 3A**). In that regard, several factors were far less abundant at the fetal than at the adult stage, suggestive of adult-specific functions. Examples are *BHLHA15*/MIST1, the master regulator of the secretory program and secretory cell architecture (Lo et al., 2017), *KLF9*, a negative regulator of epithelial and tumor cell proliferation (Shen et al., 2014; Spörl et al., 2012), and *ZBTB16* which affects diverse signaling pathways including cell cycle, differentiation, programmed cell death and stem cell maintenance (Xiao et al., 2016).

When considering the differences among adult gland types, the SL demonstrated specific enrichment in TFs over the PAR and SM, with a 5- to 10-fold increase in transcripts for genes known to regulate mucous cell formation including *FOXA1 (Ye and Kaestner, 2009), NKX2-3* (Biben et al., 2004) and *NKX3-1 (Schneider et al., 2000)* as well as TFs involved in regulating cell differentiation including a downstream effector of NOTCH signaling, *HEY1 (Nandagopal et al., 2018)*. Moreover, we discovered that many TFs often used as markers in salivary glands are differentially expressed among adult gland types, possibly suggesting differences in activity. The above-mentioned gene *BHLHA15* (*MIST1*), which has mature-gland-specific expression, exhibits differential expression at both the mRNA (**Figure 3A**) and protein levels among mature gland types, with the SM gland showing the highest expression and the SL the lowest (**Figure 3B**). Thus, our results suggest that, rather than a few gland-specific TFs driving functional diversity, specific dosage combinations of dozens, if not hundreds, of TFs likely shape the transcriptome variation of individual adult salivary glands. These outcomes also provide new TF gene signatures for the different salivary gland types that can be utilized for numerous basic biology, pathology genetics, and diagnostic applications.

We further hypothesized that the TF expression profiles of adult gland tissues can explain the expression of salivary-gland-specific expression. It then follows that there may be regulatory elements across the genome, which are activated by specific, salivary-gland-specific TF combinations. If true, we would expect genes with salivary-gland-specific expression to cluster in the genome, around these regulatory elements. To identify genes that are expressed specifically in salivary glands, we compared transcript levels in salivary glands to those of 54 other tissues in the GTEx portal, including other epithelial organs that secrete fluids such as the pancreas, mammary tissue (non-lactating), and intestine (GTEx Consortium, 2013; GTEx Consortium et al., 2017) **(Table S5, Figure S3)**. This analysis identified 188 novel transcripts with observable gene expression in adult salivary glands but negligible expression in 53 other tissues and organs reported in the GTEx database (**Figure 3C, Table S4**).

Mapping the positions of these genes, we have identified dozens of gene clusters across the human genome that show salivary-gland-specific expression (**Figure 3D, Table S4**). Some of these, such as *SCPP*, *CST* (Dickinson et al., 2002), *BPIFA* (Zhou et al., 2006), and *PRB* gene clusters, are known to have salivary-gland-specific expression. As we rely on the tissue samples available in the GTEx dataset, some of these that are enriched in salivary glands may also be expressed in other tissues not included in the GTEx database. Regardless, we reveal for the first time specific loci harboring salivary gland-specific gene sets, offering the new opportunity of investigating the regulation of gene expression in salivary glands. Excitingly, this will enable the identification of specific regulatory elements that lead to salivary gland gene expression in a tissue- or even cell-type-specific manner.

Of the 188 genes identified, 80 are thought to be protein-coding. They include genes that encode proteins that were found in the highest concentrations in saliva (e.g., amylase, MUC7, salivary histatins, salivary proline-rich proteins, and others). However, the functions for most of the others remain unknown even in other organs and tissues. Those with known molecular functions include protein-coding genes such as zinc finger proteins (e.g., the embryonic stem cell transcription factor *ZFP42* (Kim et al., 2011), immune-related genes (e.g., *KIR3DX1*), and peptidase inhibitors (e.g., *SPINK9*). However, the functions of these genes specifically in salivary glands remain unexplored. A few of these protein-coding genes were previously reported to be specific for other organ systems not included in the GTEx database, e.g., placenta-specific protein (PLAC4). The remaining 108 genes are long non-coding RNAs that to our knowledge have not been identified in the salivary gland context. Examples of these include those with known functions e.g., LINC00273, a possible regulator of lung cancer metastasis (Jana et al., 2017; Sarkar et al., 2020), and AC092159.2, which has been suggested to play a role in metabolic processes (Hu et al., 2019) as well as others with as of yet unknown functions e.g., LINC00967. Given the multiple roles of long non-coding RNAs in other organ systems, these transcripts may play a role in controlling nuclear architecture and transcription in the nucleus as well as in modulating mRNA stability, translation, and post-translational modifications in the cytoplasm of salivary gland cells. Overall, our expanded dataset provides a novel set of transcripts that can be utilized as specific markers for the salivary glands. Furthermore, it provides now dozens of gene targets for future functional studies to better understand salivary gland homeostasis, regeneration, and function.

### Transcriptional and post-translational regulation of abundant salivary secreted proteins

In addition to defining salivary-gland-derived proteins in saliva, by combining advanced quantitative technologies such as RNAseq and mass spectrometry, we can begin to dissect observed protein level variance into contributions from transcriptional, translational, and post-translational processes, as performed previously in other systems (Liu et al., 2016). Thus, to begin to determine if saliva protein abundance is regulated at the transcriptomic, translational or post-translational level, we compared transcript levels in each glandular tissue type with protein abundances in the corresponding glandular ductal secretions available through the Human Salivary Protein Wiki, a saliva-centric, curated mass-spectrometry-based proteome database that recently became available (https://salivaryproteome.nidcr.nih.gov/**; Figure 4A, Figure S4, Table S2**). This database provides quantitative data on abundances of proteins in whole mixed saliva as well as in ductal secretions obtained from PAR and SM/SL. The proteomic profiling by mass spectrometry was able to accurately quantify 2,911 proteins (**Table S6**). We found at least 10 normalized RNAseq reads per gene in at least one of the salivary glands for 85%, 81%, and 80% of the top 100 proteins expressed in whole saliva, PAR, and SM/SL secretions, respectively. Overall, we did not find a genome-wide correlation between transcript levels in either of the glands and their corresponding protein levels in the respective glandular secretions (R^2^ < 0.1). Similarly, there was no direct correlation between glandular transcriptomic abundances and protein abundances observed in the whole saliva (R^2^ < 0.1). This suggests that the vast majority of proteins in the whole saliva are either not derived from salivary glands or that protein levels are being affected by post-transcriptional processes or post-secretory enzymatic modifications that impact protein abundance.

**Figure 4.**
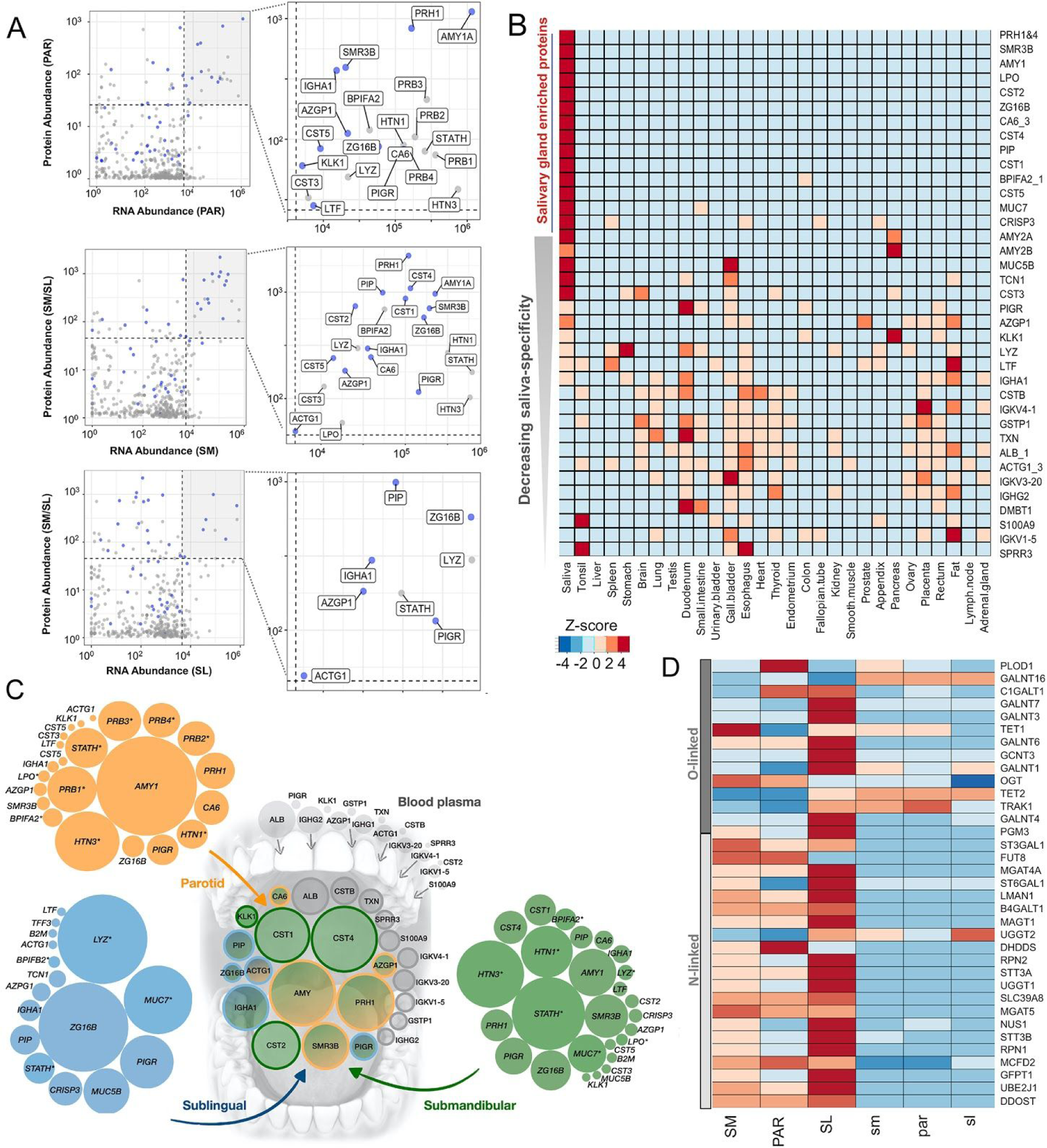
The shaping of the salivary proteome. **(A)** In order to compare transcript levels of genes expressed by the 3 types of salivary glands to protein levels in saliva, we integrated our transcriptomic data with the mass-spectrometry-derived proteome of whole saliva (Human Salivary Proteome Wiki database). Each graph represents a comparison of transcript abundances of a specific gland-type with protein abundances in that gland’s corresponding ductal saliva. The x-axis represents log_10_ normalized transcript levels, and the y-axis log_10_ normalized protein abundances. Genes showing the highest abundance (top 10%) at both the transcript and protein level are highlighted in the top-right quadrant by a grey background. The areas of the grey quadrants are shown enlarged in the right panels with the protein names indicated. Blue dots indicate genes coding for secreted proteins that are highly abundant at both the transcript and protein levels. Grey dots indicate genes that are highly abundant at the transcript level but not at the protein level. **(B)** Comparison of the most abundant proteins in human saliva (Mostly from PAR) with the proteomes of 29 human organs from the Human Protein Atlas database (Wang et al., 2019). Genes were chosen based on their expression levels from the saliva proteome and salivary gland RNAseq analysis. The data are normalized across rows for observing the comparative abundance of proteins across tissue types. The x-axis shows the tissue types and the y-axis the gene names. The genes are ordered from top to bottom based on their enrichment in salivary glands. The colors represent a z-score i.e., the deviation from the mean expression in a given row. **(C)** Glandular origins of the most abundant saliva proteins in whole mouth saliva. This schematic diagram summarizes our observations and shows the putative origins of the most abundant proteins in whole mouth saliva. The central group of circles represents the most abundant proteins detected in whole mouth saliva by quantitative mass-spectrometric analysis (data source: HSP-Wiki https://salivaryproteome.nidcr.nih.gov/). The areas of the circles approximate the observed protein abundances. Orange, green, blue, and gray colors indicate the putative origin of these proteins from the corresponding major salivary glands or from blood plasma exudate. The groups of circles on the outside represent the gene expression (RNA) levels in PAR (orange), SL (blue), and SM (green) coding for abundant salivary proteins as detected by mass spectrometry in the corresponding glandular secretions (data source: HSP-Wiki https://salivaryproteome.nidcr.nih.gov/). The areas of the circles approximate relative RNA abundances in each gland type. Note that some proteins, indicated by an asterisk (*) next to the protein name, are detected at the glandular level as secreted proteins but were not among the most abundant proteins detected in whole mouth saliva. The gray circles on top indicate the relative protein abundances in blood plasma, but only of only those proteins that were abundantly detected in whole mouth saliva. (**D)** Heatmap of expression levels of genes that are categorized by the Gene Ontology database as either “protein N-linked glycosylation” (GO:0006487) or “protein O-linked glycosylation” (GO:0006493), and are abundantly expressed (>1000 normalized gene count from DESeq) in salivary gland tissues. Note that we omitted genes listed under GO subcategory “O-glycan processing” (GO:0016266), which are mostly mucin genes. These genes encode for proteins that are targets of O-glycosylation rather than regulators. The colors indicate z-scores for each row of data. **Table S2** provides a comprehensive list of glycosylation-related genes and their gene expression in salivary glands.

Furthermore, we also noted differences between gland types in the relationship between transcript and protein abundance. As shown in **Figure 4A**, the SL appears to produce proteins found highly abundant in SM/SL ductal saliva from genes displaying lower abundance at the transcript level (<10^4^ normalized gene count by DESeq, lower panel), whereas such an outcome was not observed for the SM or PAR. However, we did find that most highly abundant proteins in ductal salivas are also generally highly expressed at the RNA level in the corresponding glandular tissues (**Figure 4A**). As shown in the upper and middle panels of **Figure 4A**, genes with the highest transcript levels (>10^4^ normalized gene count by DESeq) in the PAR and SM show also the highest abundance at the protein level. Taken together, these data indicate that transcript levels by themselves are not always sufficient to predict absolute protein levels in saliva, and that major salivary glands differ in post-transcriptional regulation.

In order to further define the origins of proteins in saliva, we next compared the most abundant proteins, ranked according to protein abundance in the human salivary proteome and according to transcript abundance by our RNAseq analysis, to the publically available mass-spectrometry-based proteomes of 29 healthy human organ tissues from the Human Protein Atlas project (Wang et al., 2019). These 29 organs included the PAR as well as other organs that secrete fluids such as the pancreas, prostate, and gallbladder (Wang et al., 2019) (**Table S5)**. Through this comparative analysis, we delineated 14 out of the top 50 secreted saliva proteins to be highly enriched in salivary glands and saliva as well as at least 5 other proteins that besides the salivary gland were found to be highly expressed in one or two of the other organs tested (**Figure 4B**). These findings which suggest that salivary glands are the organs of origin for these proteins in saliva are also supported by our comparative analysis of salivary gland mRNA to the 54 tissues/organs within the GTEx transcriptomes (**Figure S3**). However, since some of these proteins, including MUC7, LPO and PIP, are known to be present in other body fluids (not included in the GTEx transcriptome database) such as tear fluid and respiratory mucus (Jung et al., 2017; Sharma et al., 1998), they are not exclusive to salivary glands but are likely markers of multiple human exocrine organs (the pancreas being the exception). Other proteins, such as CST2, CST5, ZG16B, and SMR3B also showed little to no protein or transcript expression in other tissues or organs, including secretory mammary gland, pituitary, prostate, pancreas, and lung, and have only been detected in tear fluid (Jung et al., 2017).

We also found genes that were abundantly expressed in glandular tissues (>1000 normalized gene counts by DESeq2) but could not be detected at the protein level in salivary secretions. These genes were enriched in functions related to intracellular housekeeping processes, as well as in those typifying exocrine tissues including vesicle-mediated transport (*e.g.*, *MCFD2*), regulated exocytosis (*e.g.*, *TCN1*), and cell secretion (*e.g.*, *FURIN*) (**Table S3)**. In contrast, genes encoding proteins that were identified in saliva, but were not detected at the RNA level in glandular tissues, were enriched in functions characteristic of epithelial cells, including keratinization and cornification (*e.g.*, *KRT1, SPRR1A*). One particular protein that is abundantly found in saliva (among the top 10% abundant proteins) but is not highly expressed (bottom 10%) at the RNA level in glandular tissues, is albumin. This finding proves that the majority of albumin in the whole saliva is not derived from salivary glands but rather diffuses into whole mouth fluid from blood plasma, where it accounts for more than 50% of all plasma proteins, through epithelial leakage and gingival crevicular fluid as had been suggested earlier (Helmerhorst et al., 2018).

In addition, we noted that certain secreted proteins which were abundantly detected at the mRNA level in glandular tissues and at the protein level in ductal saliva, such as STATH, LYZ, MUC7, and HTN1, among others, were detectable by mass spectrometric analysis at much lower amounts or not at all in whole mouth saliva **(Figure 4A and Figure S4)**. This reduction or loss can be due to these proteins being proteolytically degraded once exposed to the mouth environment (Thomadaki et al., 2011), or through adsorption to oral surfaces after secretion from salivary glands. Indeed, multiple studies have demonstrated STATH, LYZ and HTN1 to be selectively adsorbed from saliva onto the enamel surface in form of the acquired pellicle, (Hannig et al., 2005; Hay, 1973; Li et al., 2004)). It is also possible that mass spectrometric analysis could not quantitatively detect certain proteins in saliva due to, for example, dense glycosylation that protects them from trypsin cleavage, or other molecular features that impede identification of unique peptides in the mass spectrometer (Thamadilok et al., 2019; Walz et al., 2006, 2009). In that regard, a recent mass-spectrometry-based proteomic analysis of healthy PAR has revealed multiple proteins including HTN1 and LYZ to be highly expressed in the glandular tissue (Wang et al., 2019), thereby supporting our prediction of protein loss after secretion from the gland.

Overall, integrating our glandular RNAseq and mass-spectrometry-derived protein abundance data, we were able for the first time to parse out the origins of proteins present in human saliva. In other words, we were able to determine which proteins in whole saliva are intrinsically produced by the salivary glands and which are derived from other sources, such as epithelial cells or from plasma leakage in the mouth. Our current understanding of these connections, as based on our own transcriptome data and on proteome data, is schematically summarized in **Figure 4C**.

As stated above, some of the most abundant proteins in saliva are heavily glycosylated, among them most prominently the densely O-glycosylated salivary mucins MUC5B and MUC7, as well as other high-molecular-weight glycoproteins including salivary agglutinin/gp340/DMBT-1, proline-rich glycoprotein, secretory immunoglobulin A, and others (Oppenheim et al., 2007). Indeed, glycosylation of salivary proteins is a major determinant of multiple aspects of oral function, from mucus layer formation to microbial binding (Cross and Ruhl, 2018). Thus, we specifically investigated whether glycosylation programs were similar or different between gland types. To test this, we analyzed the expression patterns of genes that regulate O-linked and those that regulate N-linked glycosylation, as annotated by the Gene Ontology Database (**Table S4**). We found that, indeed, each salivary gland type expresses a unique repertoire of transcripts for genes that regulate glycosylation.

Focusing on the most abundantly expressed glycosylation-related genes, it became clear that the SL shows dramatically increased expression of multiple GalNAc transferase genes (GALNTs). These enzymes are important for the initiation of O-glycosylation, a hallmark feature of mucin proteins abundantly present in salivary gland secretions. This finding is novel and makes sense biologically, given that the SL produces the major proportion of highly glycosylated mucin proteins in human saliva. Furthermore, we identified here for the first time the gene repertoire that the SL utilizes to produce its glycosylated products. It is also worth emphasizing the magnitude of this difference among gland types. For example, *GALNT12* is expressed ∼100-fold higher in SL tissue than in the other glands. We also discovered a specific GALNT enzyme that was highly specific to the SM (*GALNT13*). GALNT genes have been shown to be non-redundant in both animals and humans, and thus likely have very specialized roles in catalyzing different types of glycosylation (Bennett et al., 2012; Narimatsu et al., 2019). Overall, our results highlight the differences in expression patterns of glycosylation regulators among salivary gland types, setting the stage for future studies dissecting the functional basis generating the salivary glycome. This will become particularly important from a biomedical perspective, as the salivary glycome forms an interface with the oral microbiome (Cross and Ruhl, 2018), and abnormalities in glycosylation are now discussed as biomarkers for both Sjögren’s syndrome and oral cancers (Chaudhury et al., 2015; Nita-Lazar et al., 2009).

### Cellular heterogeneity within gland types underlines gland-specific protein secretion

To further consolidate our findings that the most abundant proteins in saliva are directly related to RNA transcript levels and show gland-specific expression, we conducted immunofluorescent imaging using tissue sections from the three adult gland types. Although many secreted proteins have been analyzed in individual glands, there has thus far been no comparative analysis of these proteins in all salivary gland types, and it was not determined if gene expression within the glands matched protein expression. Here we found clear concordance of gland-specific expression on the protein level with RNA transcript levels for a number of genes including *STATH, AMY1*, lactoperoxidase (*LPO*), cysteine-rich secretory protein 3 (*CRISP3*), *MUC7*, and *MUC5B* (**Figure 5A**). The expression patterns of each of these proteins are tissue-specific and are concordant with previous studies analyzing individual gland types (Nielsen et al., 1996; Ruhl, 2012; Veerman et al., 2003).

**Figure 5.**
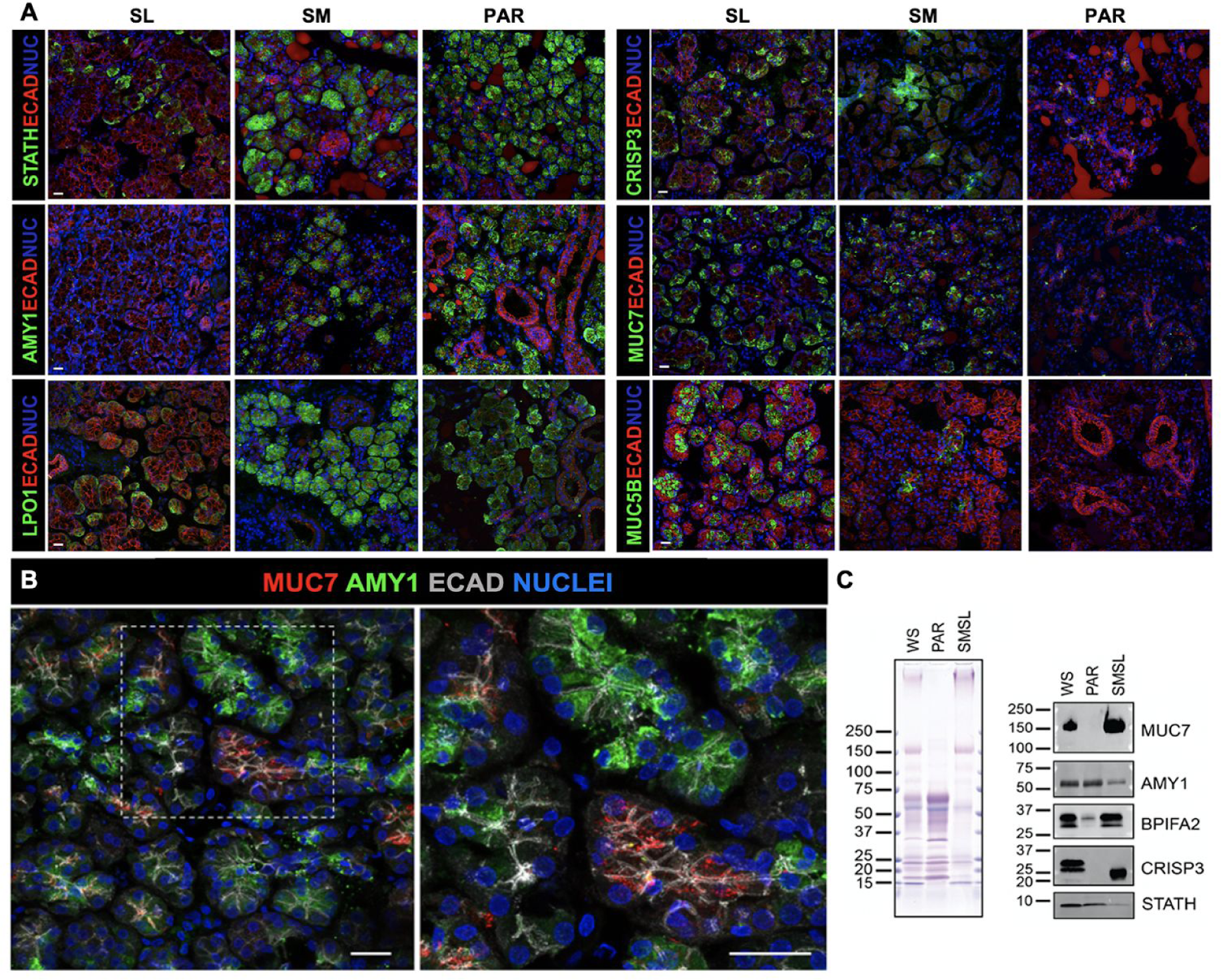
Gland- and cell-specific expression of salivary proteins. **(A)** Immunohistochemistry of glandular tissues. The SM and PAR acinar cells are highly enriched for amylase (AMY1), statherin (STATH) and lactoperoxidase (LPO) compared to the SL, consistent with these being markers of serous cells. MUC7 and CRISP3 are expressed by a subset of acinar cells of the SM and SL, with little to no expression in the PAR. MUC5B is highly expressed by the SL mucous acinar cells but not by the acinar cells of the PAR or SM, indicating it to be a marker of SL function. ECAD = E cadherin; NUC = nuclei. Scale bar = 25 µm. **(B)** MUC7 and amylase are expressed by distinct subtypes of serous acinar cells. The mature SM tissue section was immunolabeled for MUC7 (red), AMY (amylase, green) and E-cadherin (blue). (**C**) Gel electrophoretic separation of WS and glandular secretions (PAR, SMSL) followed by staining of proteins with Coomassie blue and of glycoproteins with periodic acid Schiff stain (pink bands in upper panel), and probing of transfers with antibodies against MUC7, AMY, BPIFA2, CRISP3, and STATH (lower panel).

One striking example for gland-specific expression is salivary amylase, an enzyme synthesized by serous acinar cells, that shows abundant expression at the protein level in PAR and SM glandular tissue while being virtually absent in the SL. A similar trend was found for STATH and LPO. The lower expression levels of these gene products in the SL are likely due to the lower amount of serous acinar cells in this type of glandular tissue (Amano et al., 2012). However, the near-complete absence of amylase in the SL serous acinar cells indicates that these cells are distinctly different from their counterparts in the SM and PAR. Our findings confirm the validity of using these proteins as key markers to discern SM and PAR gland-derived tissues from those of the SL.

A different example of the gland- and cell-specific expression is MUC7 which shows abundant expression at the protein level in the serous cells of the SL and, to a lesser extent, in the serous cells of the SM, while being absent from cells of PAR gland (**Figure 5A**), thus matching the RNA transcript levels from the respective glandular tissues (**Figure 1E**). Given this result illustrating the diversity of serous cells at the glandular level, we next asked if there was intraglandular or cellular variation in protein synthesis, and pursued this question by immunostaining for amylase and MUC7. Strikingly, we found MUC7 to be enriched in subsets of serous acinar cells that are deficient in amylase and vice versa (**Figure 5B**). This novel observation suggests that serous cells within the SM exist as distinct populations. Indeed, recent single-cell RNA sequencing of murine parotid salivary glands indicated a significant degree of acinar cell heterogeneity (Oyelakin et al., 2019). Thus, human acinar cells may also be heterogeneous with respect to saliva protein expression, thereby increasing the diversity of cellular origin.

In addition to serous acinar cells exhibiting heterogeneity in the synthesis of secretory proteins, we also discovered that the different gland types utilize different cell lineages for the synthesis of the same specific saliva protein. As shown in **Figure 5A**, we found that protein expression of CRISP3 paralleled that of MUC7 in being abundant in the acinar cells of the SL. However, in the SM, which expressed lower transcript levels of *CRISP3* compared to the SL, CRISP3 protein could be located only in a few acinar cells but was instead predominantly found within cells of the intercalated ducts. A similar expression pattern has been observed previously for CRISP3 protein in the murine lacrimal gland (i.e., acinar and duct cells are CRISP3+), but has not been reported for human tissues (Reddy et al., 2008). Thus, our data newly suggest that distinct human glandular cell types can synthesize the same saliva proteins, further expanding the cellular origins of these molecules.

To prove whether what we observed at the gland level by immunohistochemistry manifests itself at the protein level in salivary secretions, we conducted gel electrophoretic separation and western blot analysis of glandular ductal secretions for AMY1, MUC7, CRISP3, BPIFA2/SPLUNC2, and STATH (**Figure 5C).** As revealed by Coomassie blue and periodic acid Schiff stain, the combined secretions of SM and SL (SM/SL) glands showed strikingly different patterns of protein and glycoprotein bands compared to the PAR secretion, while whole mixed saliva showed a combination of both. Western blot analysis supported our above findings that abundant saliva proteins are expressed in a gland-specific manner and are regulated at the level of transcription. AMY was predominantly expressed in PAR secretion, with less, although still substantial levels, in SM/SL secretions, whereas MUC7 was highly expressed in SM/SL secretion and undetectable in PAR ductal saliva. These results are consistent with previous reports (Merritt et al., 1973; Thamadilok et al., 2016; Veerman et al., 1996; Walz et al., 2009) as well as with our immunofluorescent analysis of salivary gland tissue (**Fig. 5, panels A and B**). We also confirmed BPIFA2, a glycosylated protein that may function in the innate immune response, to be expressed in whole saliva (Bingle et al., 2009), and we further found it to be enriched in the SM/SL sample, and weakly expressed in PAR secretion, supporting our transcriptome-based evidence that this protein is predominantly derived from the SM.

A doublet of bands for CRISP3 was detected in whole saliva, as had been reported previously (Udby et al., 2002). Here we show for the first time that the origin for the CRISP3 protein is restricted to SM/SL secretions with no protein detectable in PAR ductal saliva, thus matching our immunofluorescent analysis (**Fig. 5A**). Interestingly, the CRISP3 band in SM/SL ductal secretion migrates at a lower molecular weight range than the double bands detected in whole mouth saliva. This outcome suggests that post-secretion enzymatic processing may have occurred, likely resulting in the alteration of CRISP3 sialylation by oral bacterial sialidases, which leads to a loss of negatively charged sialic acid moieties and, thus, retarded mobility of the protein in the electrophoretic field (Udby et al., 2002; Zhou et al., 2016). We also found STATH to be present in both PAR and SM/SL ductal secretions with higher abundance in PAR saliva (Gibbins et al., 2014; Proctor et al., 2005). STATH was also abundantly detected in the WS sample run on this gel. It has to be noted though that utmost precaution was taken during sample handling and preparation to minimize proteolytic degradation. When other samples, even of the same donor individual, were probed for STATH, only a faint band or no band at all could be detected in WS (data not shown) showing that enzymatic degradation affecting particular this component can easily occur as was described earlier (Helmerhorst et al., 2010; Thomadaki et al., 2011). Overall, our combined immunohistochemical and immunoblotting data correlate well with RNA expression levels and, thus, for the first time intimately link the fields of human salivary gland and saliva protein research.

## CONCLUSION

Our analysis of the transcriptomes of mature and fetal salivary gland tissues identified hundreds of genes that together define mature salivary glands as specialized secretory organs. We also found that fetal glands, despite the late stage of development and the glands being anatomically distinct from each other, could not be distinguished based on their transcriptional profiles. This indicates that the developmental differentiation of glandular function and functional specialization of the three mature gland types occur relatively late during fetal development. These findings pave the way for future studies dissecting mechanisms of regulation of transcriptome during glandular development and will have significant implications for *de novo* organ generation.

Our results furthermore reveal for the first time the extensive level of retention of fetal genes in adult tissues in a human epithelial organ system, strongly suggesting the involvement of these genes in both developmental and homeostatic roles. Not surprisingly, the major differences between fetal and adult tissue transcriptomes manifest themselves in the upregulation of pathways facilitating secretory and immune function in mature tissue and in the enrichment of developmentally related genes in fetal glands. What was surprising, however, was the observation that hundreds of genes are retained at significant levels in mature SL tissue, while being poorly retained by the SM and PAR. Such differential gene retention likely contributes to gland-type differences, including a more fetal-like extracellular matrix of the SL.

Our results have identified hundreds of transcripts that are uniquely expressed in salivary glands as compared to other major human tissues. These genes provide a robust set for diagnostic purposes both to identify specific glandular types, and also test for deviations in expression under pathogenic conditions such as in cancer. In an attempt to further understand the regulation of these transcripts, we verified transcription factors that show differential expressions in fetal and mature glands. More interestingly, our analysis suggests that fine-scale dosage differences of hundreds of transcription factors shape the expression landscape of mature gland types. Indeed, we found dozens of clusters across the genome where multiple genes with salivary-gland-specific expression are located in proximity. It is thus plausible that these loci harbor master regulatory sequences that are bound by a specific combination of transcription factors leading to salivary-gland-specific expression. Overall, our study provides an exciting next step by having identified dozens of candidate target transcripts and transcription factors for investigating the mechanism of gland-specific expression regulation in these loci.

Our quantitative transcriptomic and proteomic comparison also allowed us to define relationships between the level of salivary gland gene transcripts and their corresponding proteins in salivary secretions. Similar to other studies exploring the relationship between high abundance proteins and mRNA levels (Liu et al., 2016), as well as quantitative studies in other secretory cell systems (Liu et al., 2016), we found that the abundance of most highly expressed saliva proteins is regulated at the transcriptional and post-translational levels. Our data also suggest differences between gland types in terms of the glycosylation machinery that likely increases protein diversity. Further, we traced the origins of abundant salivary proteins to the respective glands or to other extra-glandular tissues and organs. Collectively, our study provides a robust framework for modeling the biological make-up of a major secretory fluid as summarized in **Figure 4C**.

Last, but not least, we provide evidence about cellular heterogeneity in human salivary glands. Specifically, we were able to identify two subsets of serous acinar cells in the submandibular gland that appeared to be specialized for expressing either AMY1 or MUC7. Acinar cells have been traditionally defined in a simple binary categorization as either serous or mucous. Now, our data suggest that these cells are more heterogeneous than previously acknowledged and that subsets of these cell types are specialized to synthesize specific salivary proteins. In addition, our data revealing CRISP3 production by serous acinar cells in the SL and by ductal and serous acinar cells in the SM demonstrate that the same saliva proteins can be produced by different cell lineages. These insights have major implications for understanding the relationship between glandular secretions and protein content of these secretory fluids and consequently inform saliva diagnostics. Furthermore, our transcriptomic analysis of healthy human gland tissue has major ramifications for *de-novo* engineering of these organs as well as determining the impact of the disease on salivary gland homeostasis and function.

In future studies, it will be important to investigate how functional salivary gland tissue differentiation, transcriptional and posttranslational regulation, and intra-glandular heterogeneity play their roles in human health. We are particularly excited about opening up the possibility of asking specific questions about human evolutionary adaptations to different environmental conditions, food resources, and pathogen challenges among geographically and culturally distinct human populations with respect to their composition of saliva (Pajic et al., 2019; Perry et al., 2007; Thamadilok et al., 2019; Xu et al., 2016, 2017).

## METHODS

### Human tissue and saliva samples

Adult salivary gland tissues were collected with informed consent from patients aged 23 to 70 years by clinicians at the University of California, San Francisco Medical Center during routine surgeries (UCSF Biorepository, the institutional review body approval number 17-22669). The UCSF pathology lab deemed the investigated tissues to be healthy. For the RNAseq library preparation, we strictly sampled from core sections of the biopsy. Additionally, we took only every third 40 μm section (3-4 sections per sample) for analysis to ensure the specificity of our sampling.

Human fetal salivary glands were harvested from post-mortem fetuses obtained from elective legal abortions with the written informed consent of the patients undergoing the procedure and the approval of the Institutional Review Board at the University of California San Francisco (IRB# 10-00768). Specimens were donated anonymously at San Francisco General Hospital.

Saliva collection was performed as approved by the University at Buffalo Health Science Institutional Review Board (Study Nr. 030-505616). See **Supplementary Methods** for details on sample collection and sex distribution.

### RNA isolation and sequencing

RNA extraction, library preparation, and sequencing were conducted using standard methodologies to generate paired-end 150-bp reads that passed standard quality control steps. The resulting reads were mapped to the human transcriptome reference (hg19) from Ensembl (Zerbino et al., 2017) and quantified using Kallisto (Bray et al., 2016). Differential expression analysis was performed by DESeq2 (Love et al., 2014) from the raw reads. All downstream analysis was conducted using custom bioinformatic pipelines available at https://github.com/GokcumenLab/glabBits (under Saliva - RNAseq). The normalized gene expression levels can be found in **Table S2**. Raw RNAseq datasets were made publically available in the GEO database https://www.ncbi.nlm.nih.gov/geo/ under project name GSE143702. More detailed methodological details can be found in **Supplementary Methods**.

### Sample quality control

We reasoned that if any contamination existed in our samples, we expect it to be mainly derived from muscle tissue that these glands reside adjacent to. Thus, as a further precaution, we now searched transcripts of genes that are known to be muscle tissue-specific and not expressed in salivary glands (*MYH3, ACTA1, PAX7, MYH8, KLHL41, SGCA, MYBPH, MYOG, MYOZ1, XIRP1, XIRP2, LDB3, TNNT3, TTN, DMD, and MYH1*).

Based on this analysis, we found evidence of slight contamination in two fetal submandibular gland samples and eliminated these from the analysis reported in this study. We still provided the expression data for these two samples in **Table S2** for reproducibility purposes. Also, if contamination were a major factor, we would expect to see more variation among different samples of the same gland type due to varying levels of contamination. As a further quality control step, we screened sequenced tissue for genes encoding inflammatory markers including *IL1b*, *TNF*, IL17, *CXCL13,* and *CCL21*, and did not find any of these to be upregulated in any of the samples.

### Immunofluorescent imaging, gel electrophoresis, and immunoblotting

The immunofluorescent analysis was performed on 12-µm-thick fixed tissue sections, as previously described (Emmerson et al., 2018). Sections were imaged using a line-scanning confocal microscope (Leica Sp5). Sample preparation, gel electrophoretic separation, staining of protein and glycoprotein bands by Commassie and periodic acid Schiff stain as well as immunoblotting were performed as previously described (Heo et al., 2013; Thamadilok et al., 2019). Methodological details and a description of the antibodies used can be found in **Supplementary Methods**.

## ACKNOWLEDGMENTS

The authors would like to thank William Lau for help with retrieving the protein abundance data from the NIDCR Human Salivary Proteome Wiki, and Tasha Lea (certified pathologist assistant) and Erica Oropeza, along with The Biospecimen Resources Program (BIOS) at UCSF, for facilitating the efficient acquisition, quality control and management of biospecimens. This work was supported by the National Institute of Dental and Craniofacial Research (NIDCR) under the award numbers 1R35DE028255 (SMK) and 2R01DE019807 (SR), and by the Tobacco-Related Disease Research (TRDRP) under award #28IR-0071 (SMK).

## SUPPLEMENTARY MATERIAL

### Supplementary Tables

**Table S1. Summary information on the salivary glands used in the study**. The tissue of origin, developmental stage (mature or fetal), sex, age, and the sample size for each category. The table is embedded in **Supplementary Methods**.

**Table S2. Transcriptome data for each of the 3 major salivary gland types.** Tab 1: Directory (Readme) that includes a description of the datasets presented in this table. Tab 2: Processed transcriptome data for each of the three major salivary gland types. The non-aligned RNAseq data are available at GEO https://www.ncbi.nlm.nih.gov/geo/ with the project name GSE143702. It will be published on July, 10th, 2020. Tab3: The raw transcriptome data (normalized gene read counts) for each tissue sample derived from the 3 major salivary gland types.

**Table S3. The results of the gene ontology analysis.** Tab1 includes a gene list that we used as input for GO analysis for easy replication. The specific categorizations based on retention or upregulation of gene expression corresponding to categories shown in **Figure 2.** The subsequent tabs show GO Biological Processes enrichments for differentially expressed gene groups. Only the top 30 enrichments with FDR < 0.0001 are shown. The last tab shows GO enrichment for genes that encode proteins that are abundantly found in saliva (among the top 10% in terms of protein abundance) but are not highly expressed (bottom 10%) at the RNA level in glandular tissues are defined as "Proteome > RNAseq" and vice versa. For each grouping, we provided the gene list for ease of replication.

**Table S4. Subsets of genes that are specifically discussed in the text.** Tabs1-3 include the genes that were found to be salivary-gland-specific, are designated as transcription factors or designated as glycosylation-related, respectively.

**Table S5. The tissues of origin for RNA (GTEx) and proteome (Human Protein Atlas).** The table is embedded in Supplementary Methods.

### Supplementary Figures

**Figure S1.**
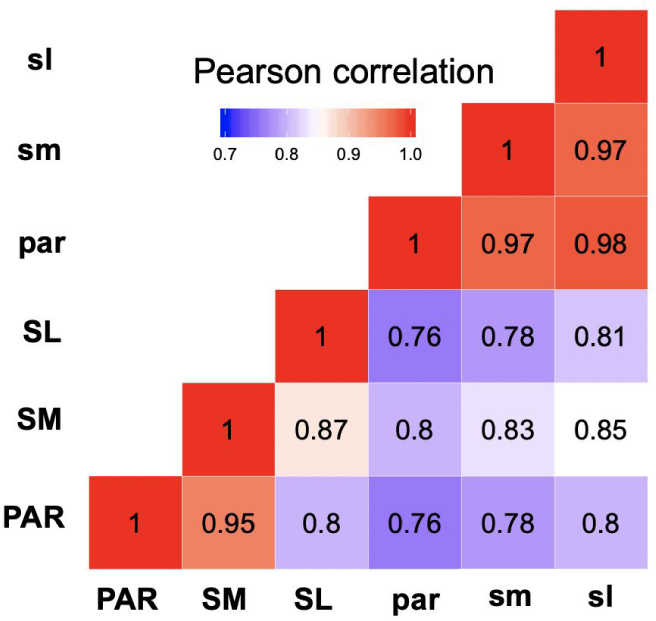
Correlation analysis of the genome-wide transcriptomes of all salivary gland samples included in this study. A heatmap was constructed by the Pearson correlation of gene expression of all salivary gland samples, with the number in each box representing the correlation coefficient among each pair compared (adult glands, upper case; fetal glands, lower case).

**Figure S2.**
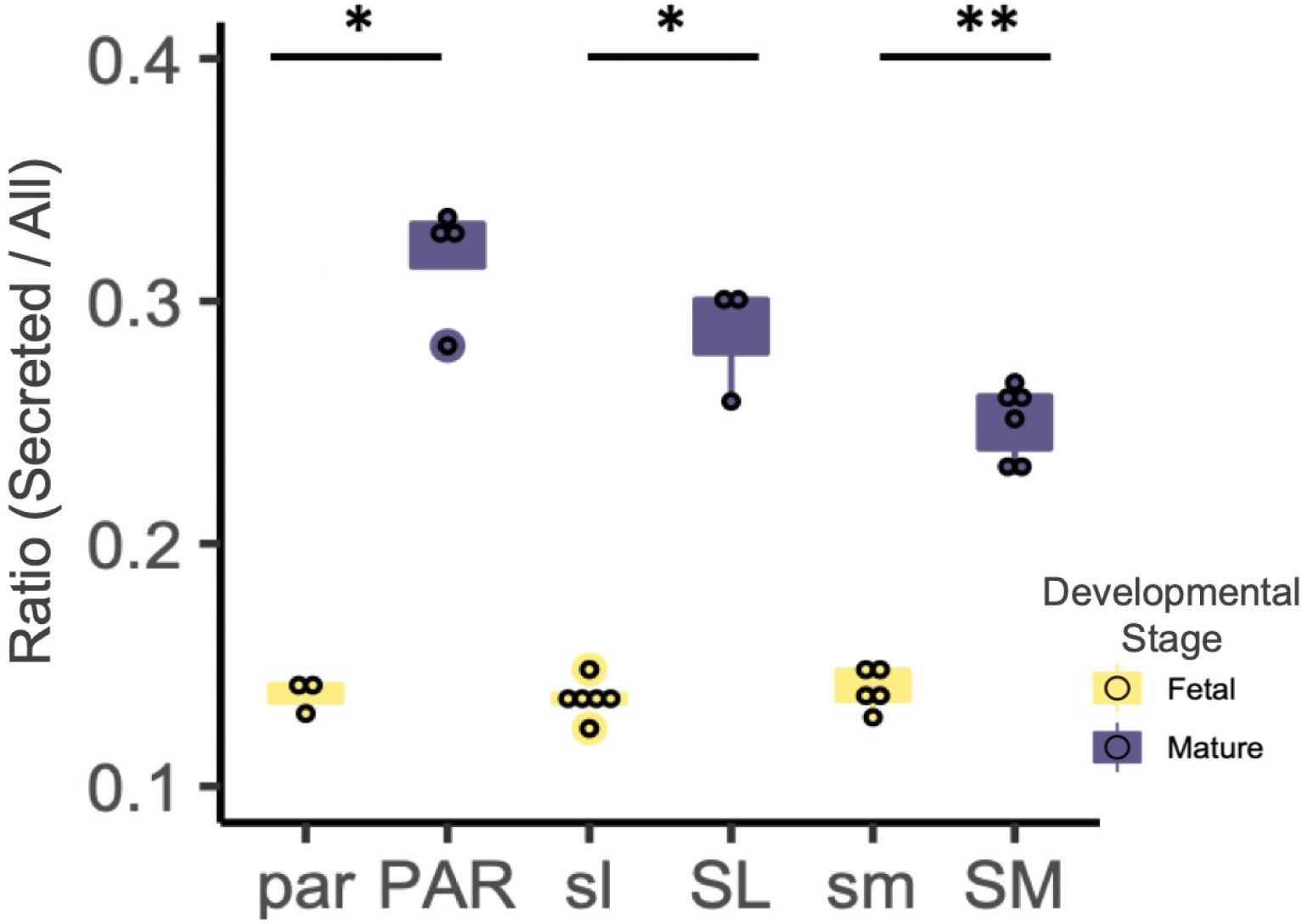
Bar plots showing the ratio of transcripts encoding for secreted proteins versus all proteins in adult and fetal salivary glands. Genes were classified based on whether they encode “secreted” proteins as per protein annotation obtained from the Human Protein Atlas (Uhlén et al., 2015). A ratio for each salivary gland was derived by dividing the sum total of all gene transcripts encoding secreted proteins by the sum total of all gene transcripts expressed. Each dot represents data from an individual salivary gland tissue sample. Mature glands show a significantly higher relative expression of genes that encode for secreted proteins than fetal glands (*p < 0.05, One-tailed Wilcoxon rank-sum test).

**Figure S3.**
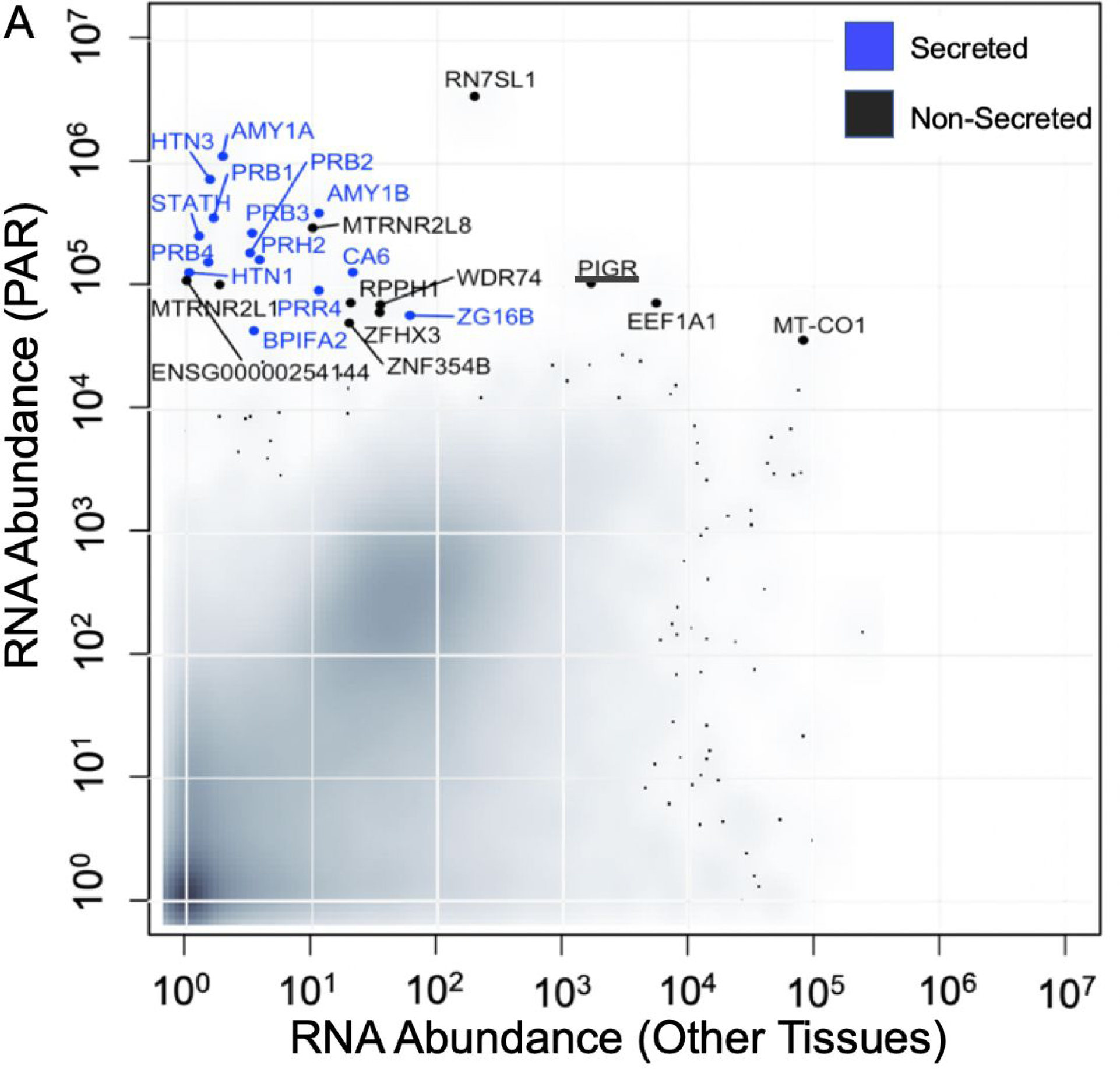
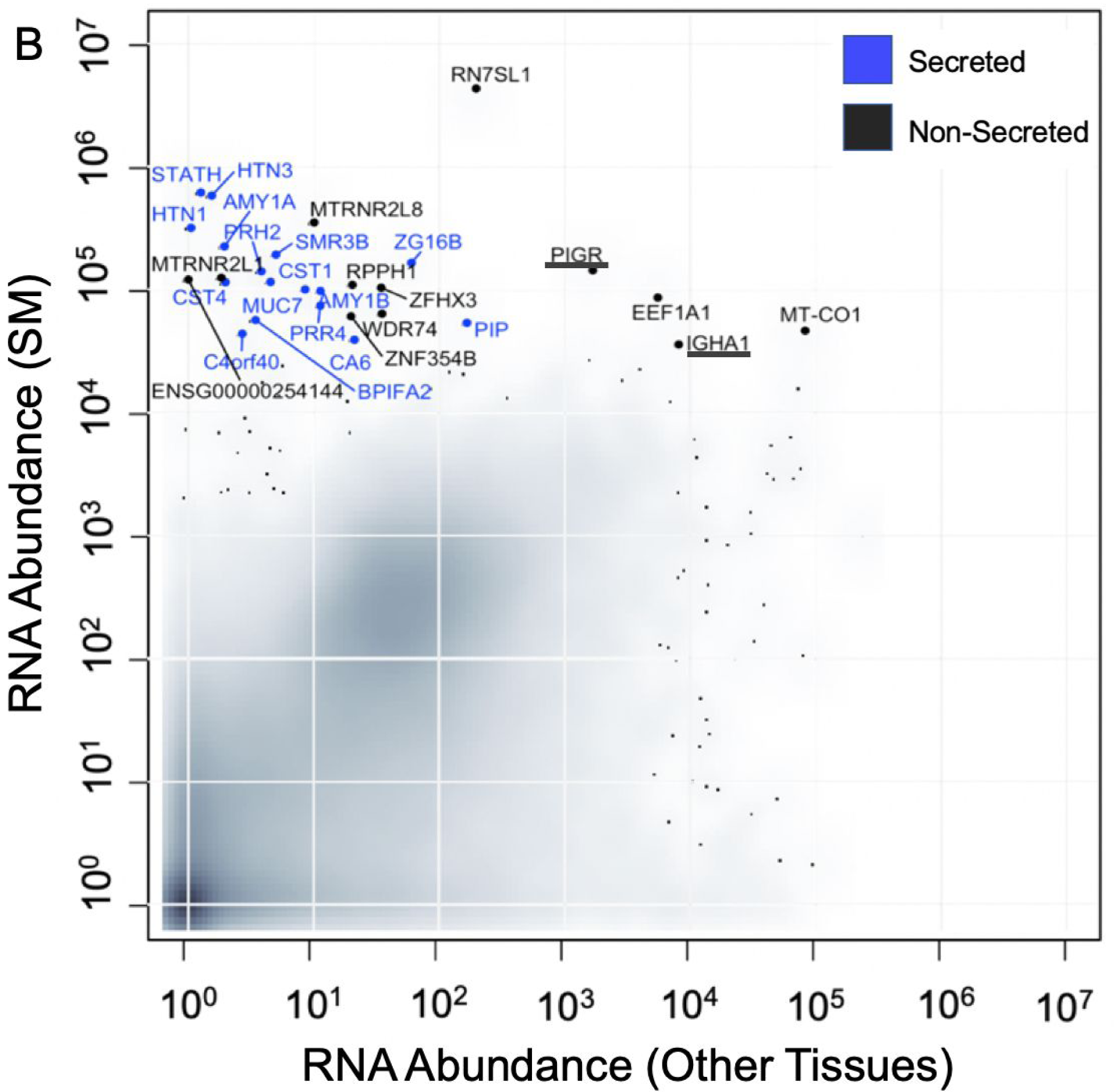
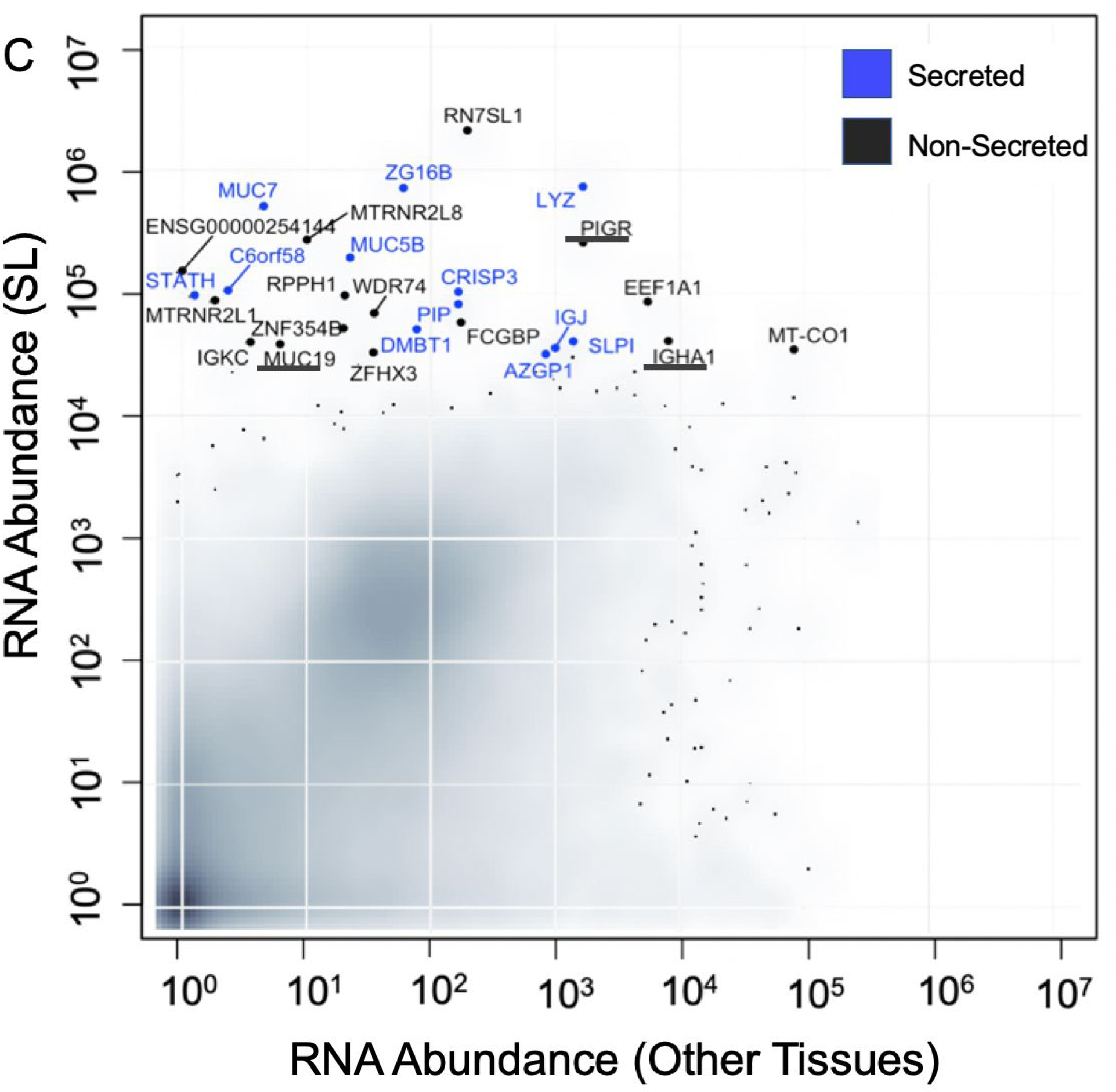
Salivary gland-specific gene expression compared to other tissues. (A-C) The y-axis shows the transcript levels of each gene (log_10_ (1+normalized read counts)) for each adult major salivary gland type. For comparative purposes, the x-axis shows the maximum gene expression levels through GTEx Portal **Table S5**. Genes coding for secreted proteins are highlighted in blue. Note that PIGR, IGHA1, and MUC19 (underlined) can occur as both membrane-bound and secreted proteins, although they were not classified as “secreted protein” in the Human Protein Atlas (see **Supplementary Methods**).

**Figure S4.**
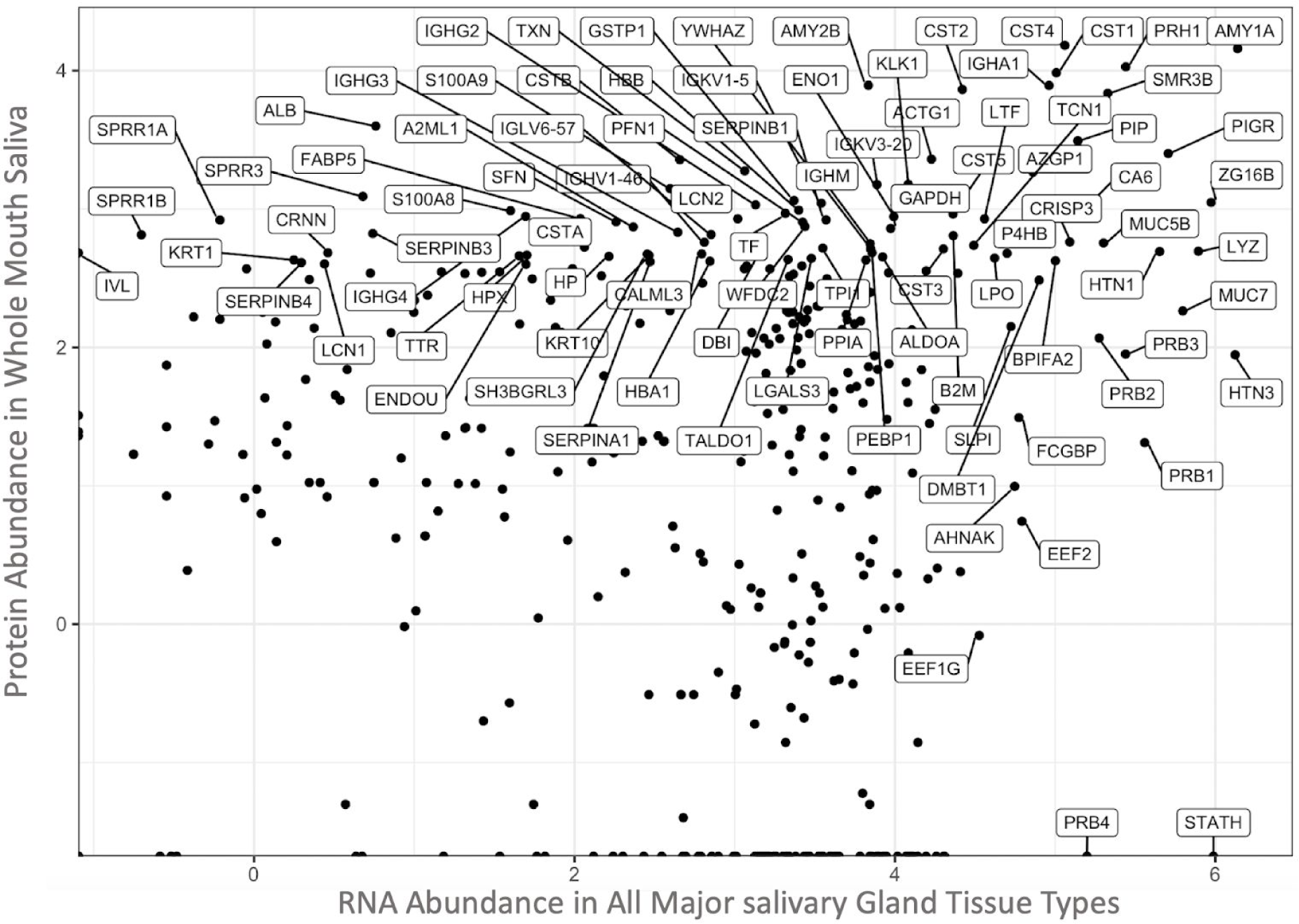
Secreted proteins in whole saliva identified by RNAseq analysis of major salivary gland tissues. The total sum of transcripts encoding for secretory proteins in all the three types of adult glands (x-axis, log_10_) was plotted against the secreted protein abundance in whole mouth saliva (x-axis, log_10_) (source: Human Salivary Proteome Wiki https://salivaryproteome.nidcr.nih.gov/). Well-known secreted proteins (*e.g.*, PIGR, AMY1A, CST4) were found highly abundant at both the mRNA and protein levels, indicating that these proteins are likely derived from the salivary glands. In contrast, non-secreted proteins that are abundant in whole saliva, such as ALB, KRT1, and SERPINB4, showed negligible transcript levels (<10 reads) in salivary gland tissue, suggesting that they originate from other tissues or organs.

**Figure S6.**
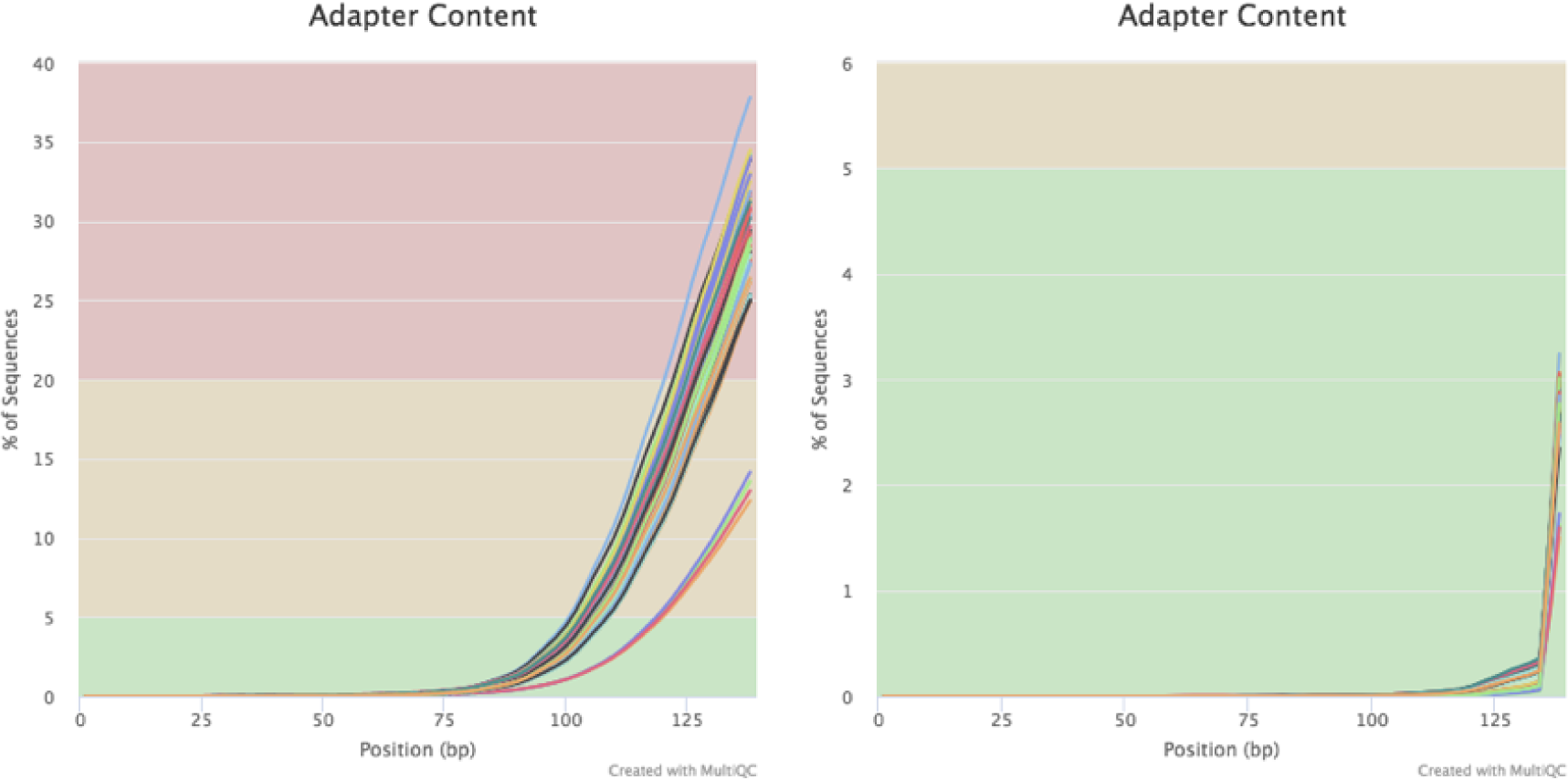
Adapter trimming of RNAseq data by using the sequence analysis tool trimmomatic. Adapter contents before (left panel) and after trimming (right panel) are shown.

### Supplementary Materials and Methods

#### Salivary gland samples

Adult human salivary gland biopsies (Table S1) were collected via the UCSF Biospecimen Resources Program (BIOS) under the institutional review body approval number 17-22669. Sample collection for the adult tissues took place during oral surgery procedures that were performed independent of this project. SM and PAR tissue samples were taken during oral surgeries from individuals suffering from cancers of the head and neck. The sample collection was limited to those patients who had not received radiotherapy, chemotherapy or immunotherapy. SL samples were derived from patients with salivary duct stones. Healthy tissue regions were identified and separated from inflamed or cancerous tissues by the UCSF pathology lab. Our immunofluorescent analysis further confirmed tissue health, as determined by cell and tissue morphology and the absence of lymphocytic infiltrates. It is unlikely but remains plausible that the disease status of the patients may have altered transcript levels. For example, it is possible that certain diseases, including oral cancers, can generate widespread inflammation, biasing our results for detecting higher levels of immune-system related genes. Since all adult sublingual glands were derived from female donors, we investigated the variation of gene expression among samples of the same gland type as well as sex-specific expression differences to test for any potential biases. We observed an extremely small variation in gene expression abundances of samples of the same gland type (**Figure 1B** and **Figure 1C**) and no sex-specific trends at the global transcriptome level (data not shown). This indicates that the differences among individual samples of the same gland type are much smaller than the gland-specific transcriptome trends we are reporting. Human fetal salivary glands were harvested from post-mortem fetuses between 22 and 24 weeks of gestation with the approval of the Institutional Review Board at the University of California San Francisco (IRB# 10-00768). Tissue was identified by location and glandular appearance. Sex was confirmed through analysis of transcript levels of male-specific genes, namely, *UTY* and *KDM5D (Staedtler et al., 2013)*.

**Table S1.** Summary information on the salivary glands used in the study.

| Number of glands and sex distribution |  |  |  |
| --- | --- | --- | --- |
| Gland type | Total number | Male | Female |
| Adult parotid | 4 | 2 | 2 |
| Adult submandibular | 6 | 3 | 3 |
| Adult sublingual | 3 | 0 | 3 |
| Fetal parotid | 3 | 2 | 1 |
| Fetal submandibular | 3 | 4 | 1 |
| Fetal sublingual | 6 | 4 | 2 |

#### Preparation of tissue samples

For RNA analysis, tissue was frozen in liquid nitrogen and stored at -80°C. RNA isolation was conducted as described previously (Emmerson et al., 2017, 2018). For immunofluorescent analysis, tissue was either flash-frozen in optimal cutting temperature compound (OCT) (Tissue-Tek) and stored at -80°C, or immediately fixed with 4% paraformaldehyde (PFA) overnight at 4°C. Fixed samples were washed with PBS, cryoprotected by immersion in a 12.5%–25% stepwise sucrose gradient, and then embedded in OCT for storage at -80°C. Tissue was sectioned (12 µm thickness) immediately before immunofluorescent analysis using a cryostat (Leica).

#### Saliva sample collection

Saliva from healthy humans was collected following the protocol approved by the University at Buffalo Human Subjects IRB board (study # 030–505616). Informed consent was obtained from all human participants. Stimulated whole saliva was collected while chewing on parafilm. Clarified whole saliva supernatant was obtained by centrifugation at 12,000 × g for 15 min at 4°C to remove particulate matter. Ductal salivary secretions were collected following the stimulation of salivary flow by application of 2% citric acid to the dorsum of the tongue. PAR saliva was collected from the orifice of Stensons’s duct using a modified Carlsen-Crittenden device, and SM/SL saliva was collected from the floor of the mouth after isolation with absorbent cotton rolls using disposable plastic Pasteur pipettes (VWR International, Radnor, PA) as previously described (Thamadilok et al., 2019). Protein concentrations in saliva samples were determined using the Pierce bicinchoninic acid (BCA) protein assay kit (Thermo Scientific, Rockland, IL), using bovine serum albumin as the standard.

#### RNA sequencing and analysis

Human adult and fetal tissue samples were mechanically homogenized using a hand-held homogenizer (Thermo Fisher Scientific) and lysed in 500 µl RNA lysis buffer (Ambion) by sonication (1 x 2-4-second pulse, Branson SFX150). RNA was isolated from 3 x 30 µm sections of human adult and fetal tissue using the RNAqueous Micro Kit (Ambion), and total RNA samples were DNase-treated (Ambion). Sample yield and integrity was analyzed using a 2100 Bioanalyzer (Agilent Technologies, Santa Clara, CA, USA). RNA sequencing was performed by standard operating procedure by GENEWIZ (https://www.genewiz.com/en) using Illumina HiSeq with a 2 x 150 bp configuration. Quality control of the obtained sequences was performed using FastQC (Wingett and Andrews, 2018) (http://www.bioinformatics.babraham.ac.uk/projects/fastqc/ Accessed 12/10/2017). The results were further reviewed by MultiQC (Ewels et al., 2016). Adaptor sequences, low-quality bases from both sides of the read (3 bases or smaller), and reads with a length smaller than 36 bp were discarded by Trimmomatic (Bolger et al., 2014) (**Figure S3**). Filtered reads were mapped to the human transcriptome reference (hg19) from Ensembl (Zerbino et al., 2017) and biomaRt (Durinck et al., 2005) and quantified using Kallisto (Bray et al., 2016). Differential expression analysis was performed by DESeq2 (Love et al., 2014), which calculates the fold-change of transcription of each gene using the Wald test and a correction for multiple hypotheses from the raw reads. We used an adjusted (i.e., multiple-hypotheses-corrected p-value of < 0.0001) to identify genes that were upregulated (fetus < adult) and downregulated (adult < fetus) during development in each type of salivary gland. Since all adult SL tissue samples were of female background, we excluded the Y chromosome from the analyses. The remaining 40,882 genes were used for subsequent analyses. The RNA abundances for 40,882 genes that we interrogated as well as comparative results are provided in Table S2. The RNA-seq data (fastq files) have been submitted to GEO https://www.ncbi.nlm.nih.gov/geo/ with the project name GSE143702. It will be published on July, 10th, 2020.

#### Immunofluorescent Analysis

Tissue section immunofluorescence analysis was performed as previously described (Emmerson et al., 2018). Frozen adult human salivary sections were fixed with 4 % PFA at room temperature (RT) for 20 min and subsequently washed in PBS followed by permeabilization with 0.5 % Triton-X in PBS for 10 min. Tissue sections were blocked with 10% donkey serum (Jackson Laboratories) and 1% BSA (Sigma-Aldrich) in 0.05% PBS-Tween 20 for 1 h at RT. Tissue sections were incubated with the following primary antibodies overnight at RT: rabbit anti-MUC7 (1:200; Sigma HPA006411), mouse anti-MUC7 (1:500; Abcam ab105466), mouse anti-MUC5B (1:500; Abcam ab105460), rat anti-E cadherin (1:300; Sigma U3254), rabbit anti-CRISP3 (1:200; Sigma HPA054392), rabbit anti-AMY1A (1:200; Sigma HPA045394), rabbit anti-AQP5 (1:400; Millipore AB3559), rabbit anti-statherin (1:500, Ruhl laboratory), and rabbit anti-LPO1 (1:200; Sigma HPA028688). Antibodies were detected by incubating samples with Cy2-, Cy3- or Cy5-conjugated secondary Fab fragments (1:300 in 0.05% PBS-Tween-20; Jackson Laboratories) for 2 hours at RT. Nuclei were detected using Hoechst 33342 (1:1000, Sigma Aldrich). Fluorescence images were obtained using a Leica Sp5 line-scanning confocal microscope.

#### SDS-PAGE and immunoblotting

Saliva samples were denatured under reducing conditions. Equal amounts of total protein (15 μg per lane for Coomassie and periodic acid Schiff stain) were subjected to separation by SDS-PAGE using 8-16% gradient Tris-glycine mini gels. Only for the detection of statherin, because of its small molecular size, Tris-tricine gels were used for better resolution (Novex, Invitrogen, Carlsbad, CA). Staining for proteins and glycans with Coomassie blue and periodic acid Schiff stain was performed as previously described (Heo et al., 2013). Stained gels were imaged using a flat-bed scanner in the transparent mode (ImageScanner III, GE Healthcare). Immunoblotting was performed as previously described (Thamadilok et al., 2019) except that for electrotransfer a BioRad Trans Turbo Blot apparatus was used. Blots were probed with the following antibodies diluted in Tris-buffered saline containing 2% milk (TBS-milk): mouse monoclonal anti-human mucin 7 (MUC7) diluted 1:500 (4D2-1D7, Abcam), rabbit polyclonal anti-human alpha-amylase 1A (AMY1) diluted 1:1,000 (HPA045394, Sigma), rabbit polyclonal anti-human parotid secretory protein (SPLUNC2B, BPIFA2) diluted 1:500 (C-20, a gift from Dr. Colin Bingle at the University at Sheffield (Bingle et al., 2009), rabbit polyclonal anti-CRISP3 diluted 1:500 (HPA054392, Sigma), and polyclonal rabbit anti-human STATH (Dundee Cell Products Ltd., Dundee, UK). As secondary antibodies, Alexa Fluor 488 tagged goat anti-rabbit or anti-mouse IgG (Life Technologies) were used diluted 1:1,000 in TBS-milk. Fluorescent bands were detected using a BioRad ChemiDoc imaging system.

### Data analysis and integration of datasets

All the downstream data analysis and comparison of our dataset with publicly available databases were conducted using custom bioinformatic pipelines written on Rstudio (v1.2.1335), R(v3.5.3) and ggplot2 (Wickham, 2009). These codes were made available through https://github.com/GokcumenLab/glabBits/tree/master/saliva_RNAseq.

To identify genes that are expressed in a salivary gland-specific manner, we compared our RNAseq results from the three major salivary glands to RNAseq data available through the GTEx database that were obtained from 53 tissues collected from approximately 1,000 individuals (https://gtexportal.org/home/documentationPage#AboutSamples) (GTEx Consortium, 2013; GTEx Consortium et al., 2017) (Table S4). These organs include those that synthesize secreted proteins such as the pancreas, mammary gland, thyroid, and intestine. To define whether a gene was specific to salivary glands, we first identified among the non-salivary gland tissues or organs those in which the gene in question was highest expressed. We then compared this level of gene expression to the gene expression levels in each major salivary gland. To account for potential differences in the RNAseq datasets from the GTEx database due to different experimental platforms and bioinformatics data analysis processes, we compared relative expression level ranks by using a (log_10_ (1+normalized read counts)) transformation for each organs RNAseq dataset.

Functional categorization of genes was determined by cross-checking using Gene Ontology Resources (http://geneontology.org/) (Ashburner et al., 2000). Then, an enrichment analysis was conducted using GOrilla (Eden et al., 2009) and Shiny.Go (Ge et al., 2020) applying default settings that enable multiple hypothesis testing (Table S3). Secreted protein gene annotations were obtained from the Human Protein Atlas (https://www.proteinatlas.org/) with the query: “protein_class:Predicted secreted proteins NOT protein_class:Predicted membrane proteins” (Uhlén et al., 2015).

**Table S4.**
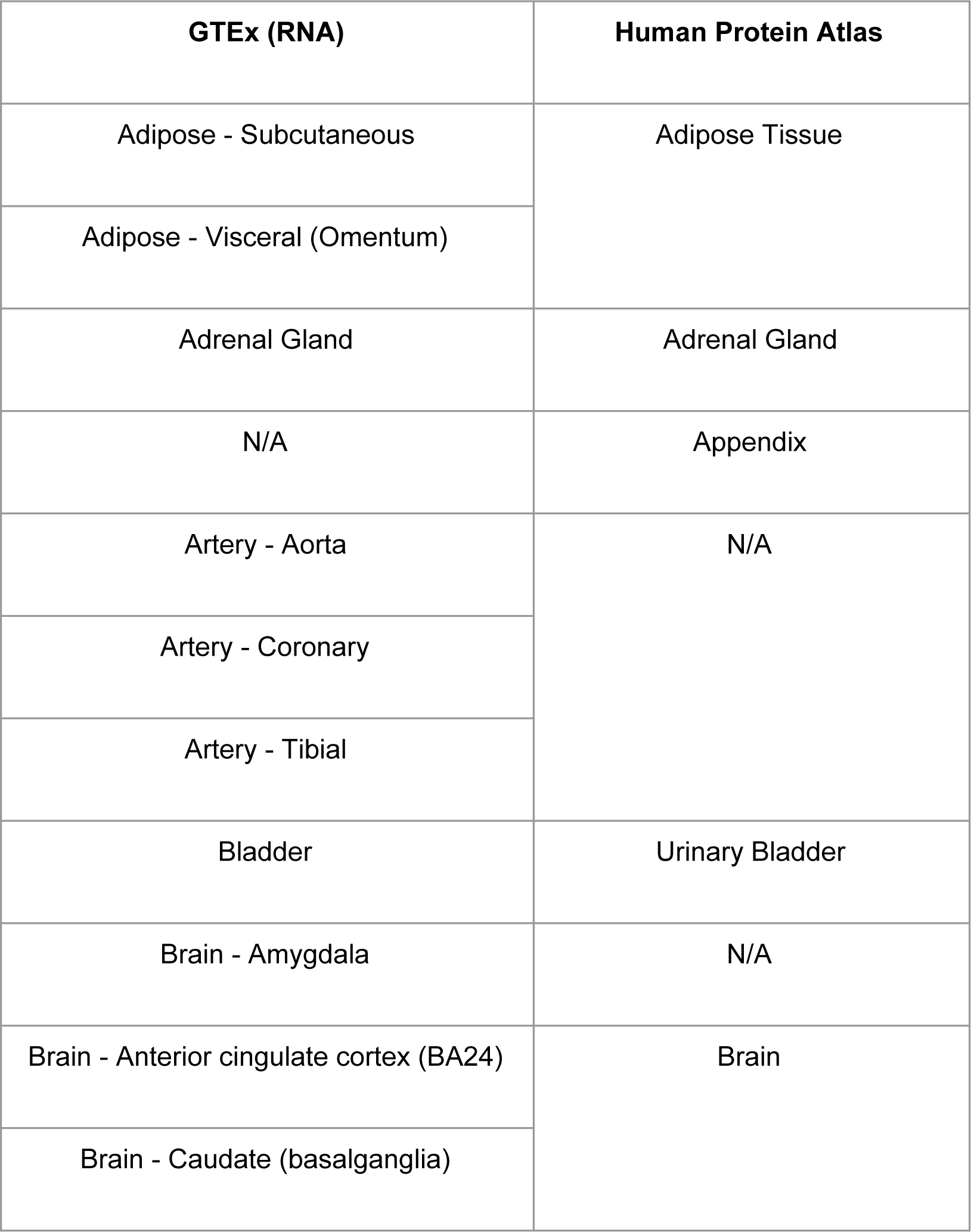
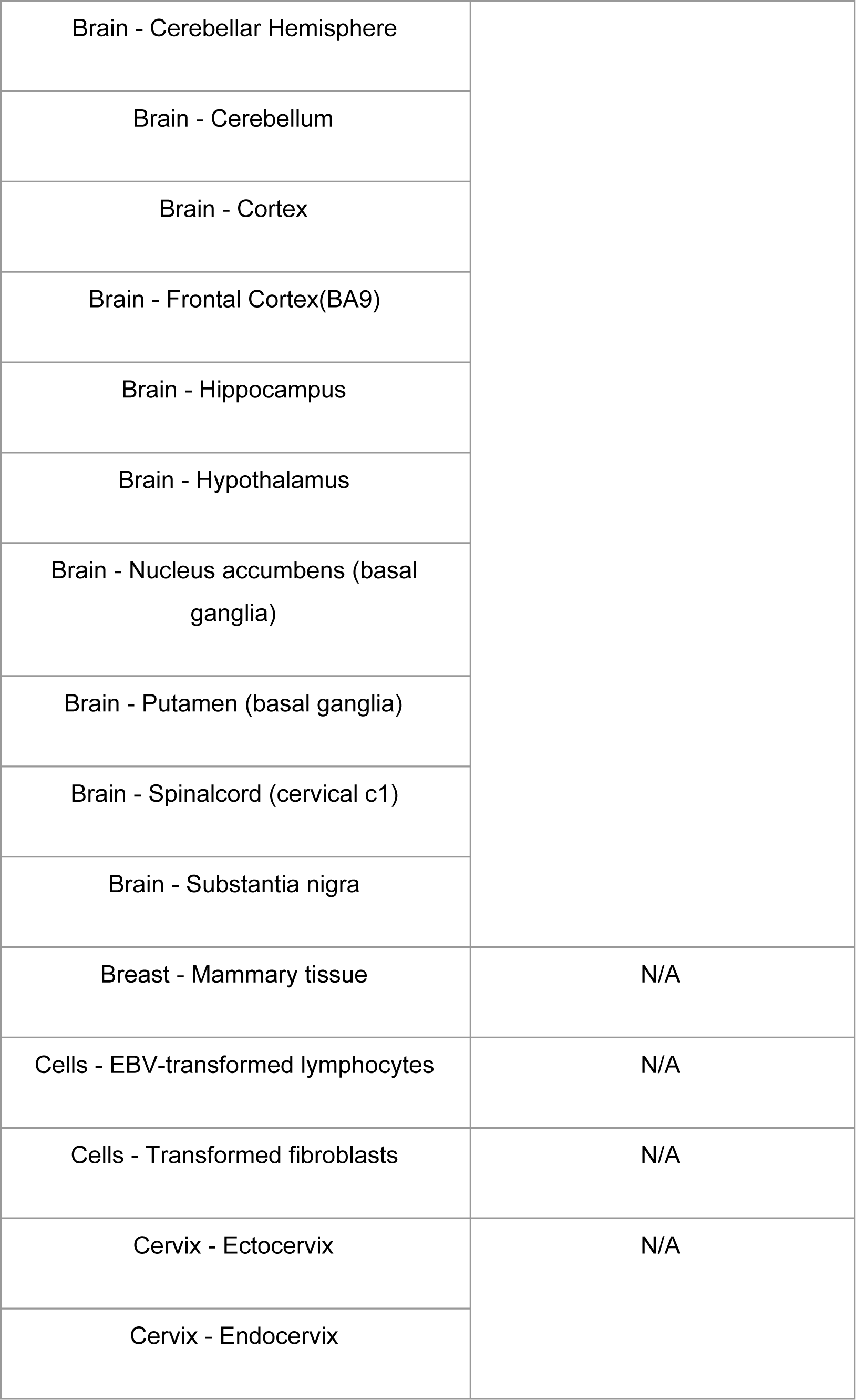
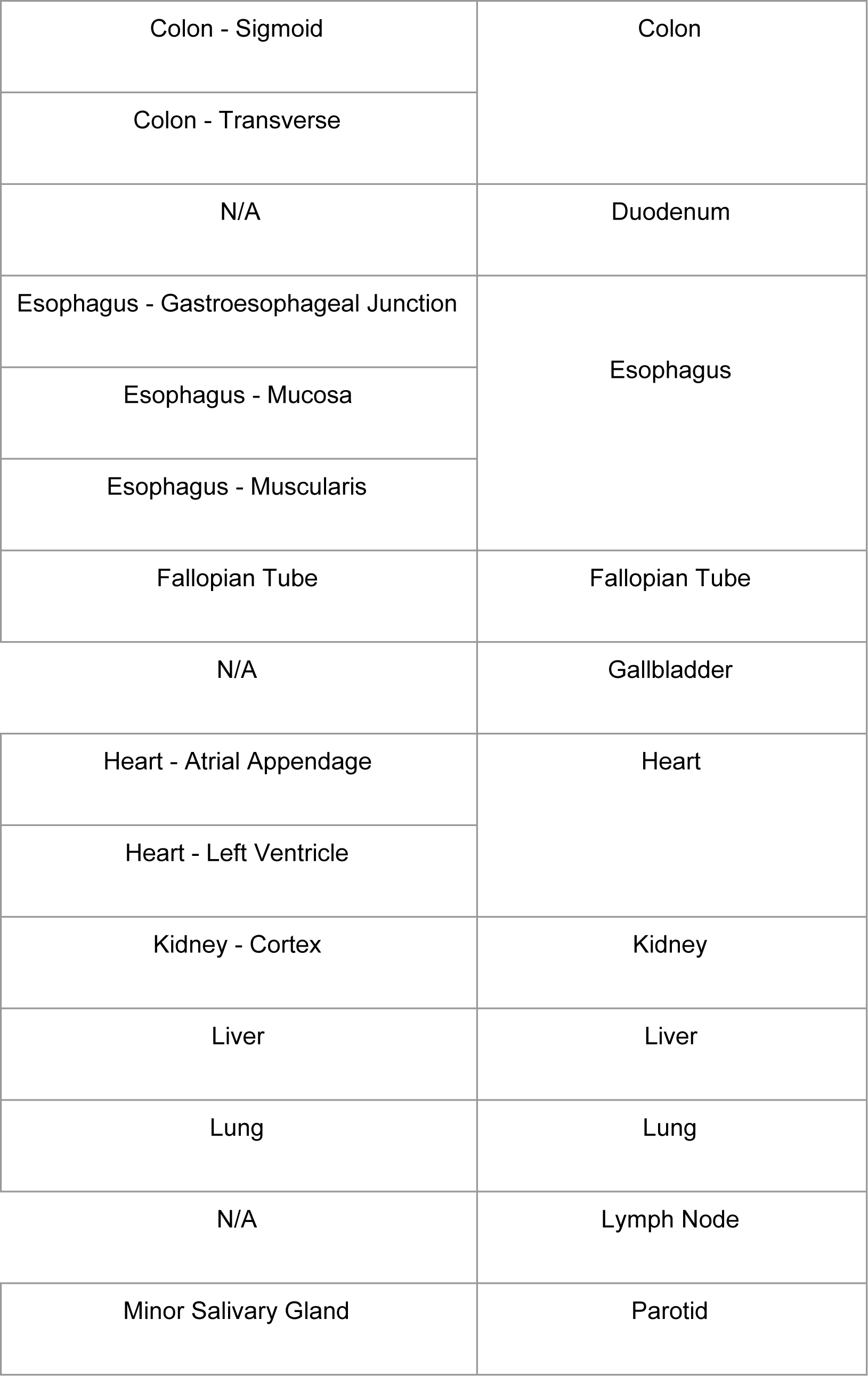
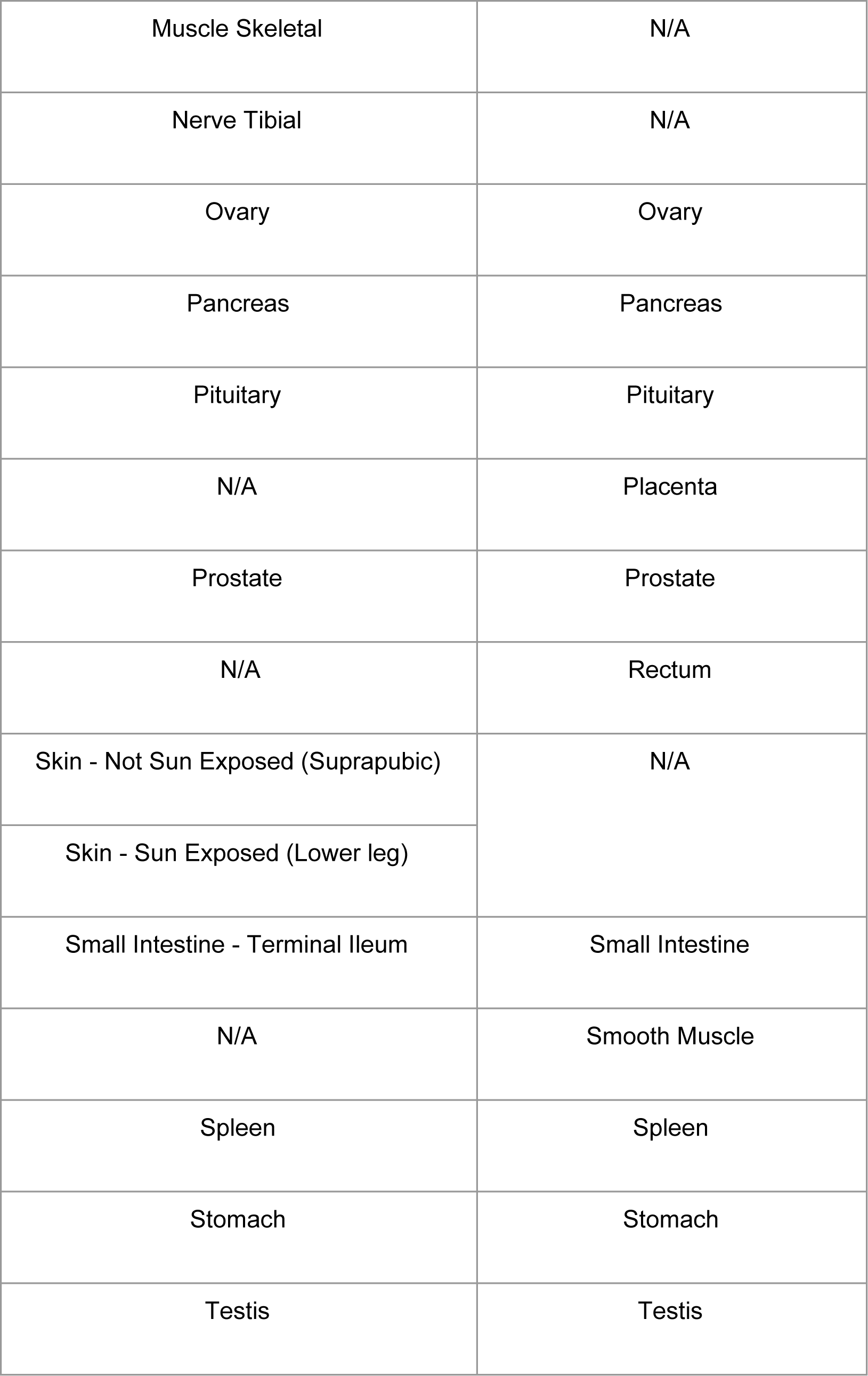
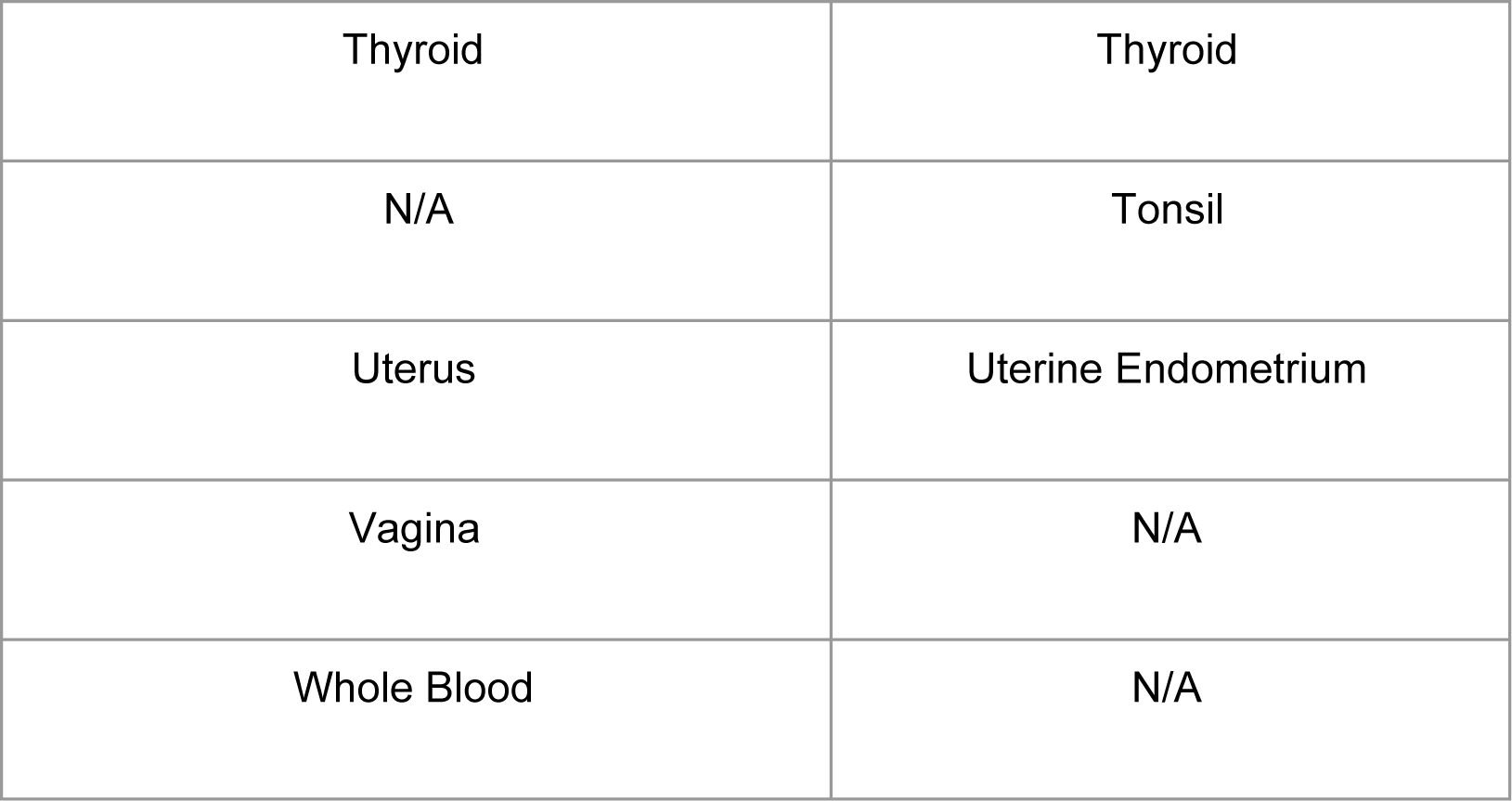
Tissues of origin for RNA (GTEx) and proteome (Human Protein Atlas **(Wang et al., 2019)**) comparison.

We used the Human Salivary Proteome Wiki (https://salivaryproteome.nidcr.nih.gov/) to obtain proteome data for whole saliva, ductal saliva from the PAR, SM, and SL, as well as from blood plasma. The database provides mass-spectrometry-based abundance data for approximately 3,000 proteins compiled from multiple studies (Murr et al., 2017), including those that focused on ductal secretions (Denny et al., 2008). Similar to the comparison approach that we used for integrating GTEx data, we used log transformation (log_10_ (1+normalized abundance)) for each dataset. In addition, we used mass-spectrometry-based protein abundances available through Human Protein Atlas from 29 tissues curated recently by Wang et al. (Wang et al., 2019) (Table S4). This dataset contains abundance information on 13,000 proteins.

